# Prior expectations about own abilities bias self-belief formation and hinder subsequent revision

**DOI:** 10.1101/2024.08.30.610443

**Authors:** Alexander Schröder, Nora Czekalla, Annalina V Mayer, Lei Zhang, David S Stolz, Christoph W Korn, Susanne Diekelmann, Finn Luebber, Frieder M Paulus, Laura Müller-Pinzler, Sören Krach

## Abstract

Self-beliefs hinge on social feedback, but their formation and revision are not solely based on new information. Biases, such as confirming initial expectations, can lead to inaccurate self-beliefs. This study uses computational modeling to explore how initial expectations and confidence affect self-belief formation and revision in novel behavioral domains. In the first session, participants developed performance self-beliefs through trial-by-trial feedback. In the second session, feedback contingencies were reversed, requiring belief revision for accurate self-beliefs. Results showed a confirmation bias in belief updating, with initial expectations being linked to biased learning during both formation and revision. Higher confidence was associated with reduced belief revision and on average, self-beliefs persisted despite the conflicting evidence. This study extends the literature on confirmation bias to learning in uncharged, novel behavioral domains. Further, it demonstrates the importance of initial expectations and associated confidence for biased self-belief formation and subsequent learning.

## Introduction

Beliefs of whether one can provide certain knowledge or perform certain tasks are crucial for everyday life. In commonplace life, we often receive feedback about work progress, personality, appearance, or abilities^1–4^. To effectively navigate daily challenges and progress toward future objectives, integrating this kind of feedback into beliefs about ourselves is pivotal^5^. Frequently, however, this feedback conflicts with how we thought about ourselves before. For example, you may consider yourself to be a funny person but later find out that your friends no longer laugh at your jokes. From the perspective of classical social psychological theory, such situations create dissonance, which people aim to resolve^6^. In this case, you have two options to resolve this dissonance: Either you reconsider your initial belief that you are a funny person (i.e., revise or adjust the self-belief), or you ignore or re-interpret your friends’ feedback and thus hold on to the initial belief. Of course, for revision to take place, an initial self-belief must already exist so that individuals can evaluate potential disconfirming feedback. In the above “joke” example, you need to have developed an existing concept of your funniness to notice that your friends’ responses are contrary to that belief. Thus, a self-belief always involves an initial expectation that has evolved in the past and describes the self-related content of the belief. Additionally, confidence in the respective expectation^7^ plays a crucial role in self-belief updating. That is the subjective “feeling of knowing” that accompanies a belief^8^. In the “joke” example, if you are very certain you are a funny person, then the self-belief of funniness would largely not be tempered. However, the exact conditions that promote revision or maintenance of self-beliefs are still unclear. The same applies to the way initial beliefs affect learning biases at different stages of learning. The present study explores the role of initial expectations and confidence in the process of self-belief formation and revision.

Previous research has repeatedly shown that individuals do not integrate information in a perfectly balanced manner to obtain the most accurate belief. Instead, this process is inherently biased^9,10^. Recent advances have benefited from computational modeling to formally quantify the underlying mechanisms of such potential biases in belief formation through feedback-driven learning. Specifically, as described by the reinforcement learning framework, beliefs are formed and updated via prediction errors (PEs), the mismatch between expectations and feedback^11–14^. Studies have emphasized the role of the valence of those PEs (positive vs. negative) in how self-beliefs are formed and how they are updated following new information^10^. For example, when confronted with information on desirable traits such as intelligence or beauty, people tend to update their beliefs more strongly after positive than negative PEs^15–17^. A similar optimism or positivity bias was found when participants were confronted with feedback on the likelihood of experiencing future adverse life events^16,18–21^.

A possible explanation for this positivity bias is that beliefs do not only help us to form accurate models of the world to maximize external outcomes but they also carry intrinsic value in themselves^22,23^. To take up the above-mentioned “joke” example, carrying the belief of being a funny person may elicit positive internal states. Individuals strive to optimize a positive affective state; thus, integrating new information is biased to support this process^22,24,25^.

In contrast, there are also occasions where it may be beneficial to assign greater weight to negative over positive feedback when updating beliefs. There is evidence showing that individuals update their beliefs more strongly after receiving worse-than-expected feedback in a performance context^1–3,26^, possibly because this feedback indicates potential for improvement^1^. Stronger integration of negative feedback was also discussed as helpful in achieving more coherence and consistency in self-beliefs^27^. Generally, several studies have shown that individuals are motivated to gain consistency in their self-beliefs by processing information in a way biased by their initial beliefs, often described as a confirmation bias^28–33^.

Especially, once beliefs are established, they become more resistant to change, which likely leads to an even stronger confirmation bias. This phenomenon has been documented across different domains of psychological research. For example, beliefs about the alleged misconduct of politicians proved to persist against correction^34^. Also, ability beliefs remain stable, even when previous feedback was invalidated as erroneous or bogus^35,36^. In the clinical context of major depression, patients also often maintain their negative prior self-beliefs and show reduced integration of new positive expectation-disconfirming information – a phenomenon labeled as cognitive immunization^37–39^. In addition, in social anxiety disorder, patients show stronger brain responses to performance feedback that corresponds to their prior negative self-assessment^40,41^. In summary, all of these biases in belief formation, namely, those motivated by self-enhancement and self-consistency, have in common that the evaluation of a belief influences how information is processed^42^.

A concept that may offer a deeper understanding of why established beliefs, at least to some extent, resist revision is that of belief confidence. Confidence guides updating behavior^43,44^. It has been shown in probabilistic learning tasks that confidence controls the weight of new evidence and how much is learned from new information^8,43^. Further, PE-based surprise responses can be amplified or dampened by confidence^44^. Those studies, however, mainly focused on confidence in decision-making tasks under uncertainty (e.g., choosing stimuli with volatile reward probabilities). Confidence is rarely measured in belief-updating studies focusing on more complex self-beliefs (like beliefs about abilities, personality traits, etc.). Nonetheless, it is reasonable to assume that once individuals have formed a self-concept, their confidence increases. Letting go of a belief that has been established over a long time is, therefore, more difficult compared to accepting new information on an almost unknown topic.

However, the exact conditions that determine the resistance of beliefs against revision remain poorly understood. Moreover, it is unclear how revision, once it does occur, relates to different motivational biases. Finally, most studies examine belief formation in contexts characterized by strong initial beliefs linked to prior individual learning histories, making it difficult to capture the conditions that support belief revision.

To bridge these gaps, the present study aimed to a) investigate the formation of epistemologically rather novel self-beliefs about the ability in an estimation task, b) characterize the revision of these beliefs in a subsequent yet separate experimental phase, and c) consider how individual differences in initial expectations and confidence impact these processes. Using a novel self-related learning task^1–3^, participants were first guided to form self-beliefs regarding unfamiliar topics (e.g., knowledge about the ability to estimate the weights of animals or heights of houses). Here, self-beliefs emerged from a neutral starting point, with low confidence in these initial beliefs (session *T1*). After self-beliefs were established in session *T1*, participants received feedback with reversed feedback contingencies to examine processes of belief revision (session *T2*). We employed computational models validated in previous work^1–3^ that incorporated trial-wise updating of the self-belief to quantify the learning mechanisms and the potential bias underlying processes of belief formation. We hypothesized that belief formation would be biased towards an individual’s initial belief, due to the valuation of initial self-beliefs and the motive to experience self-coherence^1,45^. Initial confidence in these beliefs was hypothesized to attenuate the adaption of belief-disconfirming information during belief revision.

## Results

### Measuring belief formation and belief revision

*N*=99 participants (79 female, age *M* = 22.49, SD = 2.82) completed the Learning Of Own Performance (LOOP) task^1–3^ in two consecutive sessions (12-24 hours apart, see “Methods” for details; *Figure 1b*) in the lab. The LOOP examines the formation of self-related ability beliefs based on feedback during an estimation task. In session *T1* (belief formation), participants were guided to form novel beliefs about estimation abilities through trial-by-trial feedback. In each trial, participants were first informed about the upcoming estimation category (e.g., weights of animals; *Figure 1a*). Subsequently, subjects were asked to rate how well they (*Self* condition) would perform (e.g., “I will be better than xx% of the reference group”). On the next screen, they completed an estimation task in the corresponding category and received feedback. To determine biases that are specifically self-related^1–3^, we introduced an active control condition where participants were asked to rate how well another person would perform (*Other* condition). Here, participants observed the other person’s estimation question of a different estimation category and saw the other person’s feedback.

**Figure 1.**
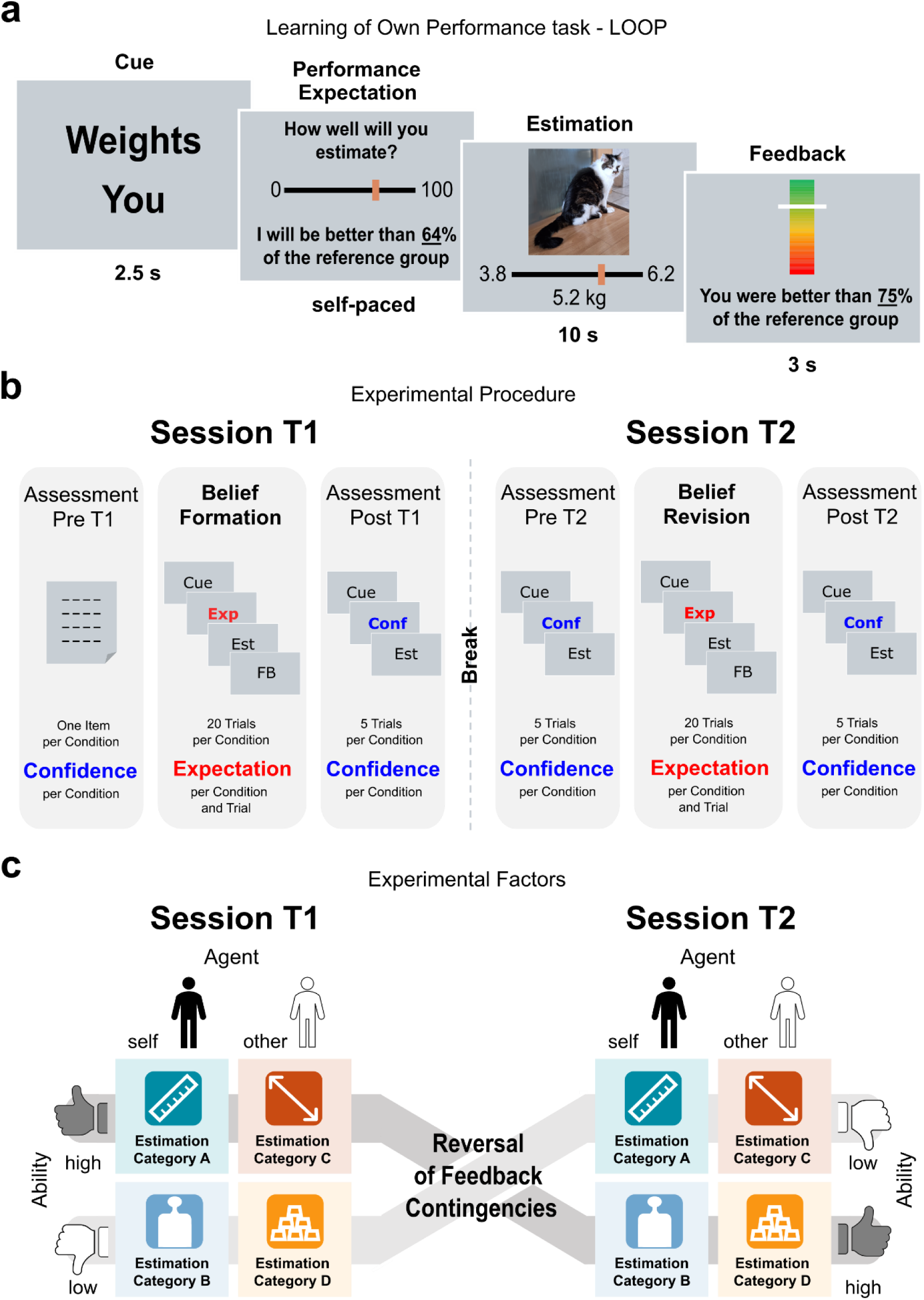
Learning of own performance (LOOP) Task and study design. **a.** Trial structure of the LOOP task. A cue prompted the estimation category (e.g., weights of animals) and agent (*Self* vs. *Other*). Participants then provided their performance expectation rating and answered the actual estimation question, followed by feedback on performance on that trial. **b.** Experimental procedure. The study was divided into two sessions. At session *T1*, confidence in the relevant ability self-beliefs was assessed in a pre-assessment. Afterward, the LOOP task was conducted to let participants form novel beliefs in the respective categories, and performance expectations were measured trial-and conditions-wise. A post-assessment was undertaken to determine the state of confidence in condition-wise self-beliefs. This was done by presenting five new estimation problems per condition, but subjects got no feedback on their performance and had to rate their confidence in how well they would perform. The mean confidence for each condition was calculated to determine the momentary level of confidence in self-beliefs. Session *T2* started with an identical assessment. Then the belief revision phase followed in which the LOOP task was done for a second time with reversed feedback contingencies. Session *T2* ended with a final assessment of belief confidence. **c.** Experimental conditions. The LOOP task was conducted on two consecutive sessions (factor *Session*: *T1* and *T2*) and additionally consisted of the factors *Ability* (*High Ability* vs. *Low Ability;* this determined whether participants were more likely to receive positive or negative feedback for the corresponding estimation category) and *Agent* (*Self* vs. *Other*). This resulted in four learning conditions that, for a given participant, were each bound to one of the four different estimation categories (heights, weights, quantities, and distances) allocated with randomized and counterbalanced order across participants. For session *T2* the feedback contingencies of the factor *Ability* were reversed. So, participants faced feedback that contradicted their ability beliefs established during session *T1*.

This was done to contrast self-related learning mechanisms with learning about others. In total, four within-subject estimation categories were distinguished: estimating the heights of buildings, the weights of animals, the number of objects, and the distance between two vehicles (pseudo-randomized across conditions). The feedback on how well they or the other person had performed was presented in comparison to an alleged reference group. Unbeknownst to the participants, the performance feedback for each trial was manipulated in such a way that there was always one estimation category associated with more positive probabilistic feedback (*High Ability* condition) and another category associated with more negative feedback (*Low Ability* condition). This design thus resulted in a total of four feedback conditions with 20 trials each (*Agent*: *Self* vs. *Other* x *Ability*: *High* vs*. Low*; *Figure 1c*; see “Methods” for details).

In session *T2* (belief revision), participants completed the same task again, but with reversed feedback contingencies, unannounced to the participants. That is, the previous *High Ability* estimation category in *T1* was now paired with *Low Ability* feedback in *T2* and vice versa *(Figure 1c)*, equally for both *Self* and *Other*. This phase thus aimed to challenge the beliefs that were established in session *T1* and thereby investigate belief revision in a targeted manner. Before and after both sessions, in additional blocks (factor *Timepoint*), participants’ confidence in their estimation-related self-beliefs was assessed (see “Methods” and *Figure 1b*).

The behavioral data was analyzed in two steps. First, model-agnostic analyses of participants’ ratings on performance expectations and confidence were conducted (detailed results can be found in the *Supplementary results*). Second, a computational modeling approach was performed to formally capture individual updating behavior on trial-by-trial basis.

### Computational modeling

To formally quantify the process of belief formation and revision we employed a computational modeling approach. We modeled the participants’ changes in expected performance through updates from prediction errors (PEs) using variants of widely used reinforcement learning models based on Rescorla and Wagner^14,46^. The winning model (1, *Figure 2b*) included separate learning rates (α) for positive and negative PEs for *Self* and *Other,* which allows a valence-and agent-specific description of updating behavior (see “Methods” for further details and the whole model space). Additionally, it contained separate learning rates for the two task sessions, i.e., belief formation and belief revision. Further, a weighting factor *w* that reduced learning rates when receiving more extreme feedback (percentiles near the 0% or 100%-mark) was included. For this, each feedback percentile was mapped onto the relative probability density of the normal distribution (ND).

**Figure 2.**
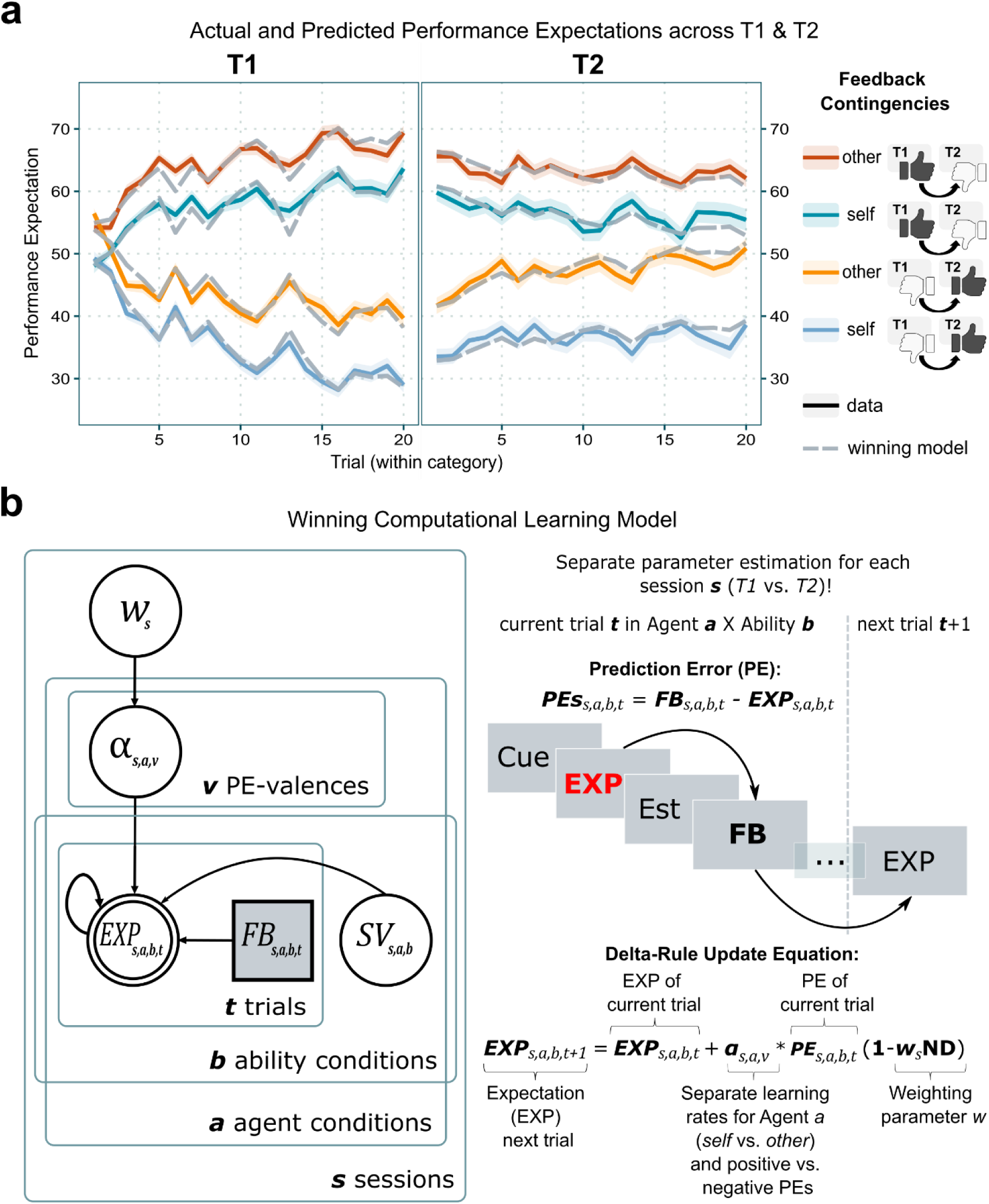
Performance expectation ratings across sessions *T1* and *T2* and details of the winning computational model. **a.** Actual mean performance expectation ratings (solid lines; shaded area: standard error) over time and mean ratings predicted by the winning model (dashed lines). During *T1*, participants adjusted their performance expectations according to the feedback given and formed distinct beliefs regarding their performance *(Self)* and the performance of another participant *(Other)* in four different domains. During *T2*, now confronted with feedback that violated the experience of the previous session, participants adjusted those beliefs to the new feedback to a lesser degree. **b.** The winning computational model. The best-performing model assumed a difference in belief updating processes for sessions *T1* and *T2*. Performance expectation (EXP) per trial and combination of the experimental factors *Agent (Self* vs. *Other)* and *Ability (high* vs. *low)* were modeled through updates from prediction errors (PEs) for each participant. PEs resulted from the differences in trial-wise expectation ratings and the feedback (FB) received. Learning rates (α) were estimated separately for *Self* and *Others* and updates following positive (PE>0) and negative (PE<0) PEs (v=valence of PE). This resulted in four learning rates per session as model parameters: α_self/PE+_, α_self/PE-_, α_other/PE+_, α_other/PE-_. This allowed a valence-and agent-specific characterization of the updating behavior across the initial belief formation phase of *T1* and the revision phase of *T2*. The winning model additionally contained a weighting parameter *w* per session, that reduced learning rates when receiving more extreme feedback (percentiles near the 0% or 100%-mark). For this, each feedback percentile was mapped onto the relative probability density of the normal distribution (ND). Lastly, the model performed best when initial beliefs (SV=starting values; the initial expectation ratings) were estimated as free parameters for each combination of the *Agent* and *Ability* conditions, resulting in four more model parameters per session.

To obtain a measure of biased learning we aggregated the estimated learning rates into a valence bias score (1) separately for *Self* and *Others* across the two sessions. A valence bias score greater than zero indicates greater feedback integration following positive PEs, whereas a score below zero indicates greater updates after negative PEs.

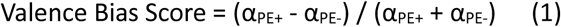

This valence bias score effectively reflects how the learning was influenced by the updates following positive versus negative PEs, as demonstrated in previous studies^1–3,47,48^.

### Self-specific negativity bias during initial belief formation (T1)

During initial belief formation, participants displayed a self-specific negativity bias during learning, replicating our earlier findings^1–3^. That is negative PEs led to higher updates as compared to positive PEs, as reflected by the learning rates. This effect was specific to the self-condition (*Figure 3a*; interaction *Agent* ∈ *PE-Valence* ∈ *Session:* β = 0.07, 95% *CI* [0.01; 0.13], *t*_(490)_ = 2.15, *p* = .032; α_Self/PE+/T1_ – α_Self/PE-/T1_: *t*_(490)_ = -6.08, *p_Tukey_* < .001). This self-related negativity bias is also reflected by the valence bias score for *T1* (*Figure 3c*, *M* = -0.14, *SD =* 0.27). In comparison, the other-related valence bias score is significantly higher (*M* = 0.11, *SD =* 0.29; *Supplementary Figure 4b*) and shows an overall positive biased updating (main effect *Agent:* β = 0.25, 95% *CI* [0.17; 0.33], *t*_(98)_ = 5.96, *p* < .001). This indicates that individuals tend to focus more on negative feedback when forming self-beliefs about their performance in new domains when the opportunity is given to improve immediately during subsequent trials^1^.

**Figure 3.**
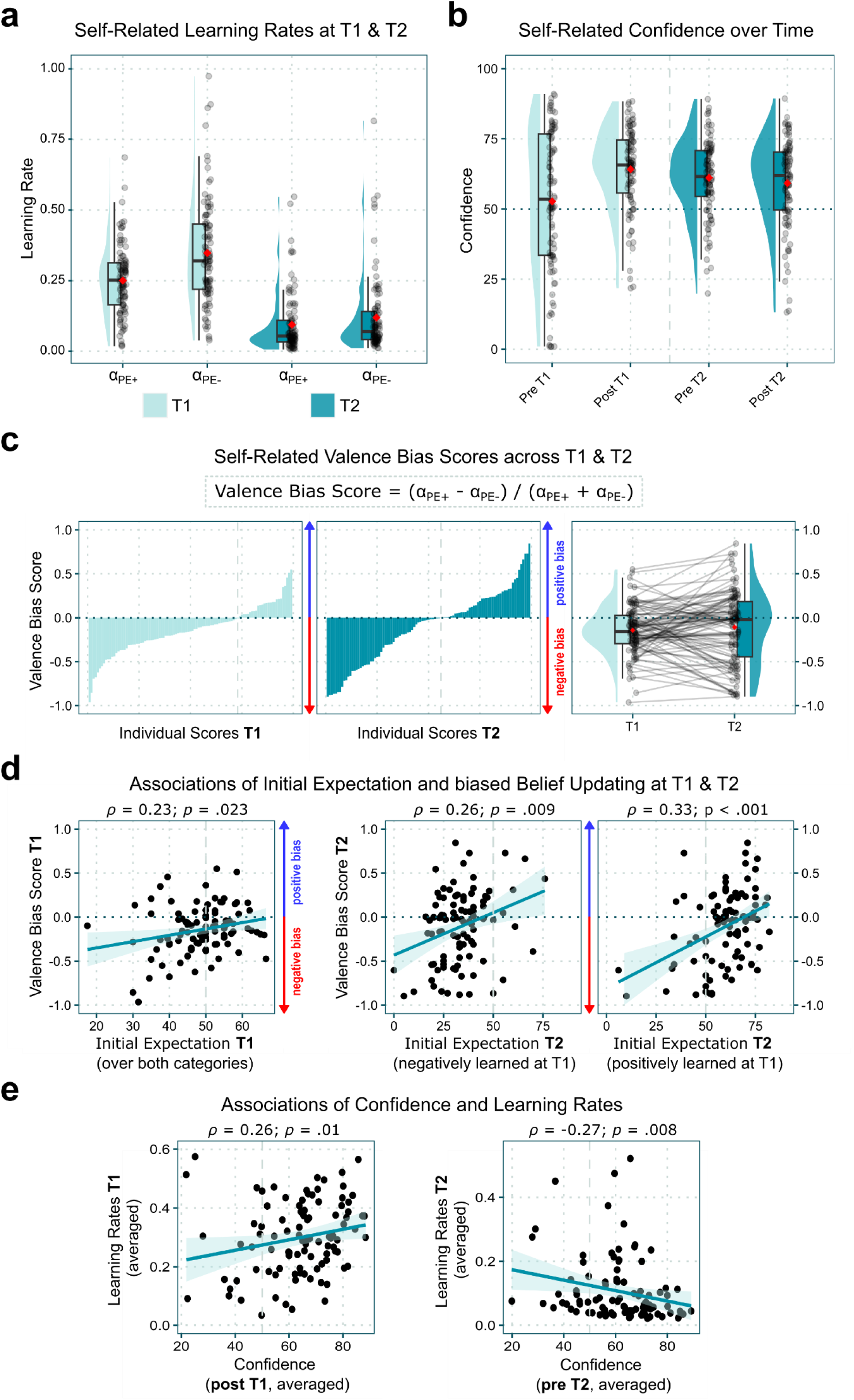
Effects of initial expectation and confidence during belief formation and revision across sessions *T1* and *T2*. **a.** Self-Related Learning Rates: Learning rates were significantly lower at *T2* than at *T1*. At *T1*, there was a bias toward greater updating after negative prediction errors (α_PE-_) compared to positive prediction errors (α_PE+_). **b.** Belief Confidence Over Time: Belief confidence increases during *T1* but only slightly decreases over *T2* when feedback contradicts the experience of *T1*. **c.** Self-related valence bias scores across the two sessions: The self-related valence bias score is calculated per session and subject by normalizing the difference in valence-based learning rates based on the sum of these learning rates. This results in a value between -1 (negative bias) and 1 (positive bias) and illustrates the subject-specific tendency for biased updating based on prediction errors. The magnitude of this value represents the discrepancy between the learning rates for positive and negative prediction errors, while it does not provide information about the extent of overall learning. The first two plots depict subject-wise scores, sorted by size and session. The third plot illustrates subject-wise changes in bias from *T1* to *T2*. **d.** Correlation plots of self-related initial expectation and the valence bias score at *T1* and *T2*. For *T1*, the initial expectations were averaged over the two ability domains since these were still unaffected by the following experimental manipulations. For both sessions, a more positive initial expectation was linked to a more positively (less negatively) biased belief updating. **e.** Correlation plots of belief confidence and learning rates (averaged over prediction error valence) across sessions. *ρ =* Spearman’s rank correlation coefficient. Boxplots in a, b, and c show median values (middle line), first and third quartiles (lower and upper lines), and mean values (red diamonds). Whiskers extend to data points within 1.5 times the interquartile range.

### Initial expectations and confidence during initial belief formation (T1)

Next, we examined how the initial performance expectation was related to belief formation during *T1* as quantified by the valence bias score. We found that a higher initial expectation was associated with more positively (resp. less negatively) biased belief formation (*Figure 3d*; correlation between initial expectation averaged over the two self-related conditions and valence bias score; *ρ* = 0.23, 95% *CI* [0.03; 0.41], *p* = .023). The same association could not be found for other-related initial expectation and valence bias score (*ρ* = 0.09, 95% *CI* [-0.11; 0.29], *p* = .359, *Supplementary Figure 5*). This suggests that self-belief formation is biased in the direction of the initial belief, even when learning in a rather new domain.

Moreover, as expected, the confidence in self-beliefs increased throughout initial belief formation (*Figure 3b*; *Timepoint post T1* vs. *pre T1:* β = 11.15, 95% *CI* [8.09; 14.22], *t*_(654)_ = 7.13, *p* < .001, see *Supplementary Note 6* for full analyses of confidence ratings). The higher the self-related learning rates during session *T1* (averaged over *PE-Valence*), the more confident participants were about their belief at the end of *T1* (*Figure 3e*; *ρ* = 0.26, 95% *CI* [0.06; 0.43], *p* = .010). When running the same analysis separated by the PE valence, results remained the same (α_Self/PE+_: *ρ* = .31, 95% *CI* [0.12; 0.48], *p* = .002; α_Self/PE-_: *ρ* = 0.22, 95% *CI* [0.02; 0.40], *p* = .031). Together, these results show that confidence in new self-beliefs changed with the rate at which new evidence in these domains was integrated.

### Diminished learning during the belief revision phase (T2)

We next examined how much beliefs were changed once they were established at *T1* but disconfirmed at *T2*. At the start of session *T2*, we found that expectations and confidence were comparable in magnitude as at the end of *T1* (see *Figures 2a* and *3b* and *Supplementary Tables 2* and *3*), showing that participants carried over their beliefs across sessions. When participants were confronted with reversed feedback contingencies during *T2*, lower learning rates for positive and negative prediction errors as compared to the initial belief formation phase were observed (main effect of *Session:* β = -0.16, 95% *CI* [-0.19; -0.12], *t*_(358.23)_ = -8.23, *p* < .001; *Figure 2a*). While learning rates for updates following positive vs. negative prediction errors did not differ significantly at *T2* (interaction *Agent* ∈ *PE-Valence* ∈ *Session:* β = 0.07, 95% *CI* [0.01; 0.13], *t*_(490)_ = 2.15, *p* = .032, α*_Self/PE+/T2_* – α*_Self/PE-/T2_*: *t*_(490)_ = -1.56, *p_Tukey_* = .405), the valence bias scores show that participants still exhibited a self-related negativity bias (*M* = -0.11, *SD =* 0.41; *Figure 3c*) and an other-related overall more positively biased updating behavior (*M* = 0.14, *SD =* 0.32; *Supplementary Figure 4b*) with no significant change in biases between the two sessions (no sig. main effect of *Session:* β = 0.03, 95% *CI* [-0.02; 0.08], *t*_(197)_ = 1.35, *p* = .178). However, due to the overall small learning rates at *T2*, it seems unlikely that these biases still had a meaningful impact at this point.

For session *T2*, again, an association between initial self-related beliefs at the beginning of *T2* and the valence bias score could be observed (*Figure 3d*; correlation analysis; former *Low Ability* category (negatively learned at *T1*): *ρ* = 0.26, 95% *CI* [0.06; 0.44], *p* = .009; former *High Ability* category: *ρ* = 0.33, 95% *CI* [0.14; 0.5], *p* < .001). This time, the same association was found for other-related initial expectations and bias (former *Low Ability* category: *ρ* = 0.21, 95% *CI* [0.01; 0.39], *p* = .038; former *High Ability* category: *ρ* = 0.29, 95% *CI* [0.1; 0.47], *p* = .003; *Supplementary Figure 5*). This indicates that the revision of self-beliefs was biased by previous beliefs in a confirmatory way: Higher expectations were linked to more positive valence bias scores. Taken together with the overall diminished learning, individuals tended to maintain their previous beliefs.

### Confidence and diminished learning during the belief revision phase (T2)

Results pointed at the impact of confidence on self-belief revision *(Figure 3e)*: the higher the participants’ initial confidence in their beliefs at *T2,* the less belief revision could be observed (quantified by the learning rates, correlation of averaged confidence *pre T2 & averaged learning rates at T2*: *ρ* = -0.27, 95% *CI* [-0.44; -0.07], *p* = .008). This showed that confidence in one’s self-beliefs might shape how strongly beliefs were maintained. Moreover, we found that participants with lower confidence in their beliefs tended to learn more from negative PEs (confidence_pre T2_ ∈ α_Self/PE+_: *ρ* = -0.07, 95% *CI* [-0.26; 0.13], *p* = .504; confidence_pre T2_ ∈ α_Self/PE-_: *ρ* = -0.32, 95% *CI* [-0.49; -0.13], *p* < .001; confidence_post T2_ ∈ α_Self/PE+_: *ρ* = -0.07, 95% *CI* [-0.26; 0.13], *p* = .518; confidence_post T2_ ∈ α_Self/PE-_: *ρ* = -0.30, 95% *CI* [-0.48; -0.11], *p* = .002), suggesting that individuals who were less confident in their self-beliefs may be more sensitive to negative information.

## Discussion

In the present study, we investigated the role of initial expectation and confidence during the formation of novel self-beliefs and, once established, during their revision. Our findings reveal self-confirmatory updating tendencies on several levels. First, we observed that the initial expectation of estimation ability was associated with subsequent learning behavior: The lower the initial expectation, the more negatively participants processed the feedback and used this information to update self-beliefs. At the same time, while the formation of self-beliefs progressed, the confidence in these newly established self-beliefs increased. This effect was evident regardless of whether participants arrived at more positive or negative self-beliefs. Further, higher learning rates were associated with higher confidence after this initial belief formation. Second, once self-beliefs were established in the first session, they were relatively robust in the face of challenging information in the second session. Also, the more confident participants were in their beliefs before learning, the less likely they used the new information to revise them. Taken together, our participants tended to process information in a self-confirmatory way, both during the formation of a novel belief and during belief revision. Our results also show that people cling to their self-beliefs even after such a short time and when they are only established in an experimental setting.

Our results are in line with evidence from research showing that individuals are motivated to gain consistency over their self-beliefs by processing information in a biased way, often described as a confirmation bias^28–32^. However, most of this research was based on tasks examining beliefs that had already been established or were related to other subjectively charged expectations^49^. For example, if a person has a high academic self-belief, he or she may be inclined to seek out information that portrays his or her academic performance in a positive light. To further dissect belief formation and explore underlying biases, it may be worthwhile to look at stages of learning in which initial beliefs are not strongly charged and are more neutral in their expectations. In the present study, we extended the confirmation bias literature to beliefs that are novel, less charged, and initially more uncertain. This was accomplished by examining self-belief formation in novel domains, such as beliefs about one’s ability to estimate weights, heights, quantities, or distances. The advantage of using such artificially constructed beliefs is that participants were rather uncertain about their potential performances at the beginning of the experiment. This allowed us to make them form new self-beliefs in either a more positive (e.g., “I think I am good at estimating the weight of animals”) or more negative direction (e.g., “I think I am bad at estimating the heights of buildings”) by manipulating the feedback in a loop-like fashion. While we replicated our previous findings of a negativity bias during self-belief formation^1–3,50^, we found that learning was linked to the initial expectation. Even when forming rather novel beliefs, people with lower initial expectations about their estimation abilities updated their beliefs in a more negatively biased way when processing performance feedback. Likewise, people with higher expectations at the beginning of the experiment showed more positive updating (or more precisely, a less negative updating). Our findings show that learning asymmetries, such as the negativity bias found in our studies, may, to some extent, be related to self-confirming updating tendencies that become apparent when initial expectations are low.

We further observed that the participants’ confidence in their beliefs increased with task progression. Thus, our targeted feedback, in one domain more positive and in the other more negative feedback, not only led to changes in expectations but also strengthened the confidence in these beliefs during their formation. This finding is also in accord with a recent publication showing that confidence changed when prior beliefs were manipulated^51^.

The change in confidence during belief formation had direct effects on processing conflicting feedback. Once beliefs were experimentally established and participants were more confident in their abilities, participants largely ignored disconfirming evidence. Although participants learned from the feedback as indicated by learning rates greater than zero, results from the modeling revealed that learning rates were significantly lower than during the belief formation phase. This attenuated updating with increasing confidence reflects another aspect of a confirmation bias, described as a lack of response to conflicting evidence^33^. This could indicate that individuals have implicitly prioritized maintaining a sense of self-consistency over optimizing belief accuracy^22,23^ by strongly adhering to their previously formed self-beliefs.

Our finding of attenuated belief revision with increasing confidence is consistent with evidence from studies using probabilistic learning tasks, which showed that confidence guides updating behavior and learning from prediction errors^43,44^. Furthermore, we found that in the face of conflicting feedback, confidence did not decrease significantly throughout the revision session. In cases where evidence for the respective self-beliefs has already been accumulated, higher levels of confidence and the maintenance of those seem justified and, in some cases, may even be adaptive by protecting individuals against potentially erroneous updates. After all, in everyday scenarios, as important as it is to incorporate self-related information into our beliefs to navigate daily challenges^5^, it is also important to protect ourselves from updates that may result from random noise. Thus, as stated by Meyniel^44^, a confidence-weighting mechanism could be a major aspect of adaptive learning. In addition, motives like promoting self-consistency and well-being also apply here. For example, it has been argued that (over-)confidence in one’s beliefs may have emerged from the motive of promoting self-consistency^52^. Moreover, being confident in one’s beliefs may in itself promote a sense of self-consistency and thus self-serving value in a way that is independent of objective belief accuracy^24^. This may also explain why participants in our study underutilized the opportunity to revise their previously negatively learned self-beliefs in the face of much more desirable feedback. Since the respective estimation abilities do not have much impact on everyday life, high confidence in an (in)ability may be valuable enough.

Notably, one cannot completely rule out that attenuated learning during the revision phase was caused by decreasing task engagement since the participants were already familiar with it. However, model-based learning rates and trial-by-trial changes in expectation were indicative of attendance at the feedback and supported the notion that new evidence was processed. In addition, the association of confidence with reduced learning during belief revision suggests that reduced learning reflects meaningful individual variance.

This study provides several advancements in the understanding of self-belief formation. First, self-belief formation, while being a continuous process overall, is characterized by forming an ad-hoc belief with a certain confidence and consecutive biases in learning with each step of incoming information. From a methodological point of view, assessing initial belief formation and later revision may allow us to study self-belief dynamics in different contexts and domains. Second, people tend to process information in a self-confirming way, both during initial belief formation and during belief revision. Finally, people cling to their self-beliefs even when they are newly formed under experimental conditions.

These findings may have important implications in educational, political, and clinical contexts. Literature on educational psychology shows that self-beliefs and academic achievements are reciprocally linked^53,54^. Translating the present findings into this frame of reference would suggest that once students have established a certain self-belief, e.g., "I am not good at math," after initial exposure to the issue, it might be very difficult to revise such a negative self-belief in the face of evidence to the contrary, especially if they have developed a strong confidence in their self-assessment. In political contexts, the literature shows many examples of rigid and hard-to-revise beliefs^34,55^. Although the direct application of our results to this context is more complex, since political beliefs only become self-relevant through additional motives or context (e.g., through the personal consequences of political decisions or, more abstractly, through the value of being part of a group characterized by specific political beliefs), it can be observed that confidence plays an important role here, too. For example, concerning rigid beliefs that contradicted consistent scientific evidence during the COVID-19 pandemic, it was shown that those who disagreed most with the societal consensus knew less about the objectively relevant aspects but were highly confident in their beliefs^55^. Finally, our findings may have implications for research on interventions aimed at improving the revision of maladaptive beliefs in clinical conditions such as major depressive disorder. This condition is characterized by negatively biased feedback processing and negative self-beliefs^56^. This also applies to experimentally established beliefs, where symptom burden is associated with reduced learning from positive feedback in patients with depression^50^. Affected individuals also lack confidence in their abilities^57^ or to put it differently, express high confidence in their lack of abilities. This high confidence in the validity of maladaptive self-beliefs may promote cognitive immunization^37,38^ -the insensitivity to new information that would objectively allow a positive belief revision. Interventions might focus on changing confidence in beliefs, similar to challenging metacognitive beliefs^58^. Without this, one should bear in mind that when challenging beliefs by exposing individuals to positive feedback, they might not be able to integrate it effectively^50^. A stronger focus on disconnecting new learning experiences from prior beliefs by promoting adaptive information processing might be a way to enhance an adaptive belief revision.

## Conclusion

In conclusion, this study demonstrates confirmatory self-belief updating on different levels: Lower initial expectations about own abilities are linked to a more negative bias. This underlines the importance of initial expectations for how we learn about the self and maintain a sense of consistency. Further, high confidence in these self-beliefs hinders their revision in the face of conflicting information. Together, these results extend research on confirmation bias to novel and initially less charged self-beliefs. Clinically, this is relevant for two reasons: first, the emergence of various mental disorders strongly depends on how self-related beliefs are formed. Second, to the extent that psychotherapy relies on revising established maladaptive beliefs, it is important to further our understanding of rigid self-beliefs and how they can be changed. More broadly, when we are interested in how humans integrate information about themselves, it is important to consider which priors individuals have and how confident they are. This may help gain further insights into the individual and contextual aspects, biases, motives, and affective components that play a role in self-belief formation and revision.

## Methods

### Participants

A total of 102 participants (79 females) between the ages of 18 and 31 years (*M* = 22.49, *SD* = 2.82) took part in two studies (see below for details). The two samples (*n_1_* = 40, *n_2_* = 62) were pooled for the analyses of this project. We excluded three individuals who did not believe the cover story of the task or did not complete the task attentively until the end (study 1: one exclusion, study 2: two exclusions), as indicated by self-reported tiredness and almost no variance in their expectation ratings. Thus, the final sample consisted of 99 participants, all of which were recruited from the University Campus of Lübeck, were fluent in German, and had either normal or corrected-to-normal vision. All participants provided written informed consent. The research received approval from the ethics committee at the University of Lübeck (reference number: AZ 21-217). It adhered to the ethical standards outlined by the American Psychological Association.

### The Learning of Own Performance task (LOOP)

#### Task explanation

The LOOP task allows participants to gradually acquire knowledge about their own or another individual’s perceived ability to estimate various unfamiliar attributes or properties, such as the height of houses or the weight of animals. This task has been previously introduced and validated through a series of behavioral and neurocomputational studies^1–3^. In this study, all participants were invited in pairs to take part in an experiment allegedly related to cognitive estimation. In cases where a second participant was not available, a trained confederate staged one. Participants were informed that they would take turns (factor *Agent*) either performing the estimation task themselves *(Self)* or observing the other person perform it *(Other)*. During each trial, participants received manipulated performance feedback in two distinct estimation categories, one associated with more positive feedback (70% positive PEs and 30% negative PEs) and the other with more negative feedback (30% positive PEs and 70% negative PEs). These conditions were assigned to the properties pseudo-randomly (e.g., height of houses = *High Ability* condition and weight of animals = *Low Ability* condition, or vice versa), and the estimation categories were counterbalanced between *Ability* and *Agent* conditions *(Self* vs. *Other)*. This resulted in four feedback conditions, each consisting of 20 trials (combinations of *Agent* and *Ability* conditions: *Self-High, Self-Low, Other-High, Other-Low*). Performance feedback was provided after each estimation trial, indicating the participant’s or the other person’s current estimation accuracy compared to an alleged previously tested reference group of 350 university students. This feedback was conveyed in terms of percentiles (e.g., "You are better than 94% of the reference participants"). The feedback was determined by a sequence of fixed prediction errors (PEs) rather than fixed feedback values. Designing the feedback sequence this way kept PEs relatively independent of participants’ expectations and, therefore, maintained sufficient statistical variability in feedback sequences between participants. More specifically, feedback was based on the current ability belief of each participant, which was calculated as the average of the last five performance expectation ratings per category, plus the respective predefined PE for each trial. Before participants provided their first performance expectation ratings this average was set to 50%. This ensured a relatively equal distribution of negative and positive PEs across conditions (see *Supplementary Table 11*). At the start of each trial, a cue indicated the estimation category and the agent responsible for the trial (e.g., height and "you" or "Ben"). Participants were then asked to provide their performance expectation rating in a self-paced fashion on a scale using the same percentiles as used for feedback. To motivate honest responses, participants were told that accurate expected performance ratings would be rewarded with up to 6 cents per trial. In other words, the closer their expectation rating matched their actual feedback percentile, the more money they would receive. Thus, reward incentives were tied to performance prediction accuracy rather than estimation performance. Following each performance expectation rating, participants had 10 seconds to answer the estimation question, with continuous response scales available for selecting plausible answers. Next, feedback was presented for 3 seconds (“You were better than 75% of the reference group.”). The task was executed using MATLAB Release 2019b and the Psychophysics Toolbox^59,60^.

#### Repeated measurements using the LOOP task

Participants completed the LOOP task in two sessions (factor *Session*: *T1* and *T2*). For study 1, these sessions were 24 hours apart. This first study was used to test the feasibility of a repeated study design that uses the LOOP task. For the second study, this time interval was shortened to 12h, and one group of participants completed session *T1* in the morning and *T2* in the evening of the same day, while a second group completed *T1* in the evening and *T2* in the morning of the consecutive day. To address potential confounds, in additional tests, *Group (wake vs. sleep)* was included as a factor in the linear mixed models that captured basic behavioral effects (see *Supplementary Notes 2, 4, and 6*), and no relevant effects could be found. To assess belief revision, in both studies, participants at *T2* received estimation questions from the same categories they had dealt with at *T1* but with inversed feedback contingencies (e.g., the estimation category “height of buildings” switched from conditions *Self* and *High Ability* at *T1* to *Self* and *Low Ability* at *T2* and the category “weight of animals” from *Self* and *Low Ability* at *T1* to *Self* and *High Ability* at *T2*; same for contingencies of condition *Other*). So, participants experienced feedback that was inconsistent with *T1*’s learning experience, which allowed them to revise their established ability beliefs. The sequence of prediction errors remained the same for both sessions.

#### Assessment of pre-and post-expectation and confidence

The very first and very last expectation ratings per condition and session of the LOOP task were used to assess *pre-* and *post*-*T1*/ *T2* expectations. Additionally, participants rated their belief confidence. Pre-and post-task confidence ratings were assessed four times (factor *Timepoint*): At the beginning and the end of the initial learning session *T1* (*pre-T1* and *post-T1*) and the beginning and the end of the revision session *T2* (*pre-T2* and *post-T2*). Participants were confronted with five new estimation stimuli from the same estimation categories as in the main task. They were asked to rate their confidence in their performance expectation rating, that is the confidence in their ability to predict their (or the other person’s) performance accurately (“How confident are you in this assessment?”). The scale ranged from zero (not certain at all) to 100 (very certain). At this stage, no feedback was provided to assess belief confidence over multiple trials without updating in between. The mean values of these confidence ratings per condition and assessment were used for further analyses. The very first assessment of belief confidence (*pre-T1*) was done with a simple query and without the presentation of specific stimuli as we wanted to capture an initial assessment that was unaffected by later aspects of the task.

#### Questionnaires and debriefing

Before commencing the experiment at *T1*, all participants were required to respond to a series of inquiries regarding demographic data. After completing the study at *T2*, participants underwent an interview that included assessments of their self-beliefs, were debriefed about the cover story, and were compensated for their time before departing. The entire procedure took approximately 3 hours – 1.5 hours for each session.

### Statistical analysis

#### Behavioral analysis

All analyses were conducted in R version 4.1.2^61^, unless stated otherwise. We initially conducted model-agnostic analyses on the participants’ performance expectation ratings and confidence ratings. To do so, we calculated several linear mixed-effects models using the *lme4* package^62^. For each dependent variable (expectation ratings at *T1*, expectation ratings at *T2*, confidence ratings) the maximal feasible random effects structures that still lead to model convergence were assessed using the *buildmer* package^63^. For expectation ratings, this stepwise procedure started from models with *buildmer* random effects structures for each session that included the *Ability* condition *(High Ability* vs. *Low Ability)* and *Agent* condition *(Self* vs. *Other)* as factorial variables and *Trial* (20 trials) as continuous predictor and that treated the Intercept, *Ability* condition, *Agent* condition, and *Trial* as both fixed and random effects. For belief confidence, it started from a similar maximal model that included the *Ability* condition (*High Ability* vs. *Low Ability*) and the *Timepoint* (*pre T1, post T1, pre T2, post T2)* as factorial variables and as both fixed and random effects. After finding the maximal feasible random effects structures, these random effect structures were included in intercept-only models, and the respective fixed effects were added iteratively. The models were estimated using Maximum Likelihood (ML). The final models to be reported were then re-estimated using Restricted Maximum Likelihood (REML) as suggested by Meteyard and Davies^64^. To further explore specific effects or interactions of the final linear mixed models, where appropriate, pairwise comparisons of estimated marginal means were conducted using the package *emmeans*^65^. See *Supplementary Notes 2* and *6* for further details and results.

#### Computational modeling

After the model-agnostic analyses, we delved into formally quantifying the dynamic changes in self-beliefs, specifically the performance expectation ratings, by employing PE delta-rule update equations, under the reinforcement learning framework with the Rescorla-Wagner model^46^. The basic equation used for these learning models is as follows (2 and 3; where “EXP” represents performance expectation rating, “FB” stands for feedback, “PE” denotes prediction error, “t” denotes the trial, and “α” signifies the learning rate):

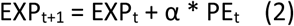

while

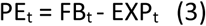

Within our model space (see *Supplementary Figure 1* for the complete model space), three primary models were explored, each with distinct assumptions regarding biased updating behaviors during self-belief formation. The simplest learning model, the Unity Model, employed a single learning rate for all conditions for each participant, thus assuming no learning biases. The Ability Model, the second model, incorporated separate learning rates for each *Ability* condition, indicating context-specific learning. The Valence Model, the third model, introduced separate learning rates for positive PEs (α_PE+_) and negative PEs (α_PE-_) across both ability conditions, suggesting that the valence (positive or negative) of the PE influences self-belief formation. Furthermore, learning rates were estimated separately for *Self* vs. *Others* across the *Agent* condition. Additionally, we tested additional model variates of these three primary models: The first assumed that individual learning rates do not vary between initial belief formation and revision (α*_T1_ =* α*_T2_*). The second assumed that individual learning rates do differ between the two sessions (α*_T1_ ≠* α*_T2_*). For the model variate that expected differing learning rates for *T1* and *T2*, two additional models were tested: The Expected PEs Model separated learning rates across positive and negative PEs and over expected and non-expected PEs. The rationale behind this model was that participants quickly learn to expect PEs of a specific valence depending on the estimation category (e.g., positive PEs in the estimation category in which mostly positive PEs were experienced), especially during session *T2*, when a belief is already established. Since the Valence Model has been the winning model in our previous studies^1,3^, specifically an extended version of the Valence Model in our most recent study that used the LOOP task^2^, this Extended Valence Model was also included. This model builds upon the Valence Model by introducing an additional parameter *w* as a weighting factor to reduce learning rates when approaching the extremes of the feedback scale (percentiles near 0% or 100%). The relative probability density of the normal distribution (ND) was associated with each feedback percentile value (see *Supplementary Figure 2*). This addon assumes that more extreme feedback would be perceived to be less likely and, therefore, provides less information. The subject-and session-specific weighting factor *w* was employed to reflect the degree to which the relative probability density influenced the reduction in learning rates for feedback far from the mean. In addition to the learning rates, the initial beliefs about one’s performance (about the other participant’s performance, respectively) were estimated separately for *Self* and *Others* as well as for both *Ability* conditions as free parameters. Taken together, the following Extended Valence Model emerges (4), where “s” represents the session (*T1* vs. *T2*), “a” the *Agent* condition (*Self* vs. *Other*), “b” the ability condition (*High* vs. *Low*) and “v” the valence of the PE (PE+ vs. PE-):

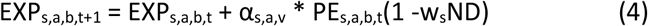

To determine whether participants’ performance expectation ratings could be better explained by PE learning compared to stable assumptions in each *Ability* condition, we also included a straightforward Mean Model with mean values for each task condition.

#### Model fitting

The RStan package^66^ for the statistical computation language R^61^ was used for model fitting with Markov Chain Monte Carlo (MCMC) sampling. For all participants, the models were fitted individually, and posterior parameter distributions were sampled. After 1000 burn-in samples 2400 samples were drawn (3400 samples in total; thinned with a factor of three) using three MCMC chains. Convergence of the MCMC chains was checked by inspecting *Ȓ* values for each parameter. Effective sample sizes *(n_eff_)* of model parameters were checked to be mostly bigger than 1500 (*n_eff_* values are estimates of the effective number of independent draws from the posterior distribution of model parameters). To summarize posterior distributions of parameters, the mean was calculated, and thus, a single value per model parameter and participant was used for the following model-based analysis.

#### Bayesian Model Selection (BMS)

To determine the winning model, the pointwise out-of-sample prediction accuracy for all models of our model space was estimated separately for all participants using leave-one-out-cross-validation (LOO; in this case, one trial-observation was left out per participant^67,68^). For this, Pareto smoothed importance sampling (PSIS) based on the log-likelihood calculated from the posterior parameter simulations was used as implemented by Vehtari and colleagues^67^. The sum of PSIS-LOO scores and the estimated shape parameters of the generalized Pareto distribution *k̑* were calculated. Only very few trials resulted in insufficient values for *k̑* and, therefore, in potentially unreliable PSIS-LOO scores (winning model *T1*: *k̑* > 0.7: 1.54%, winning model *T2*: *k̑* > 0.7: 0.39% ^67^; see *Supplementary Table 1* for all PSIS-LOO scores and % *k̑* values > 0.7). To account for group heterogeneity in the model that best describes belief updating behavior, Bayesian Model Selection (BMS) on the PSIS-LOO Scores was performed^69^ using the VBA (Variational Bayes approach) toolbox^70^ for Matlab^59^. The protected exceedance probability *(pxp)* was provided for each model. The *pxp* value indicates how likely it is that a particular model explains the data with a higher probability than all other models in the considered model space. Bayesian omnibus risk *(BOR)* was calculated, which is defined as the probability of choosing a null hypothesis (in this case, a posterior probability that shows that model frequencies are all equal in a given model space) over an alternative hypothesis^69^. In *Supplementary Table 1* PSIS-LOO difference scores of the winning model in contrast to the other models of the model space are provided. These can be interpreted as a simple “fixed-effect” model comparison^67,68^. For our data, the comparisons of PSIS-LOO difference scores were generally comparable to the results of the BMS. To check if the data predicted by the winning model could capture the variance in performance expectation ratings of each participant, posterior predictive checks^14,71^ were conducted by predicting the time course of expectation ratings for each participant and by comparing these predictions against the actual behavioral data for each session. *Figure 2a* visually confirms the model’s ability to capture the observed data for sessions *T1* and *T2*. In addition, we tested whether a model-agnostic analysis of the predicted data would capture the same effects as the analysis of the actual performance expectation ratings (see *Supplementary Note 3).* Since differences in belief updating for initial belief formation (*T1*) and belief revision (*T2*) seemed plausible, BMS was conducted separately for *T1* and *T2* to test whether different models fit best at different stages of belief updating.

#### Statistical analysis of modeling and behavioral data

Learning rates of the winning models of each session were analyzed within-subject using Linear mixed-effects models in an approach identical to that used to analyze expectation and confidence ratings as described in “model-agnostic analyses”. We started from a maximal model that included the factors *Agent (Self* vs. *Other), PE-Valence (PE+* vs. *PE-),* and *Session* (*T1* vs. *T2*) as factorial variables and as both fixed and random effects (refer to *Supplementary Note 4* for detailed results). The maximal feasible random effects structure was assessed and included in an intercept-only model. Fixed effects were then added iteratively. Further, normalized valence bias scores were calculated to associate self-or other-related learning biases with (initial) expectations and confidence (valence bias score = (α_Self/PE+_ -α_Self/PE-_) / (α_Self/PE+_ + α_Self/PE-_))^1,2,47,48^. The valence bias scores for both sessions and conditions were then analyzed in the same way as learning rates, starting from a maximal model including the factors *Agent (Self* vs. *Other)* and *Session (T1* vs. *T2).* Spearman correlations were conducted between valence bias scores, individual learning rates, (initial) expectation ratings, and confidence. All statistical tests were performed two-sided.

## Data and code availability

The code and behavioral data for the statistical analyses are available at https://osf.io/8x4vp/ (DOI 10.17605/OSF.IO/8X4VP). *RStan* scripts to estimate the computational models are available from the corresponding author upon reasonable request.

## Acknowledgments

We want to thank Alica Steinert, Sophia von Krauss, Jovana Lehmann-Grube, Finn Moritz Borcherding, Laura Rosenbusch, and Clara Scheffel for their invaluable assistance with data collection. This research was funded by the German Research Foundation (Temporary Positions for Principal Investigators: MU 4373/1-1; MU 4373/1-3; Sachbeihilfe KR 3803/11-1; Sachbeihilfe KR 3803/41-1) and the Department of Medicine at the University of Lübeck, CS08-2023.

## Author contributions

A.S., N.C., L.M.P., S.D., and S.K. designed the research. A.S. and N.C. acquired the data. A.S. analyzed the data and prepared the manuscript. A.S., N.C., A.V.M., L.Z., D.S., C.W.K., S.D., F.L., F.M.P., and S.K. discussed the data analyses and interpretation of the results and reviewed and edited the manuscript.

## Declaration of interests

The authors declare no competing interests.

## Notes

https://osf.io/8x4vp/

### Competing Interest Statement

The authors have declared no competing interest.

### Summary of Updates

- Title, abstract and conclusion revised

