## Supplementary Material for "Prior expectations about own abilities bias self-belief formation and hinder subsequent revision"

### Supplementary Figures

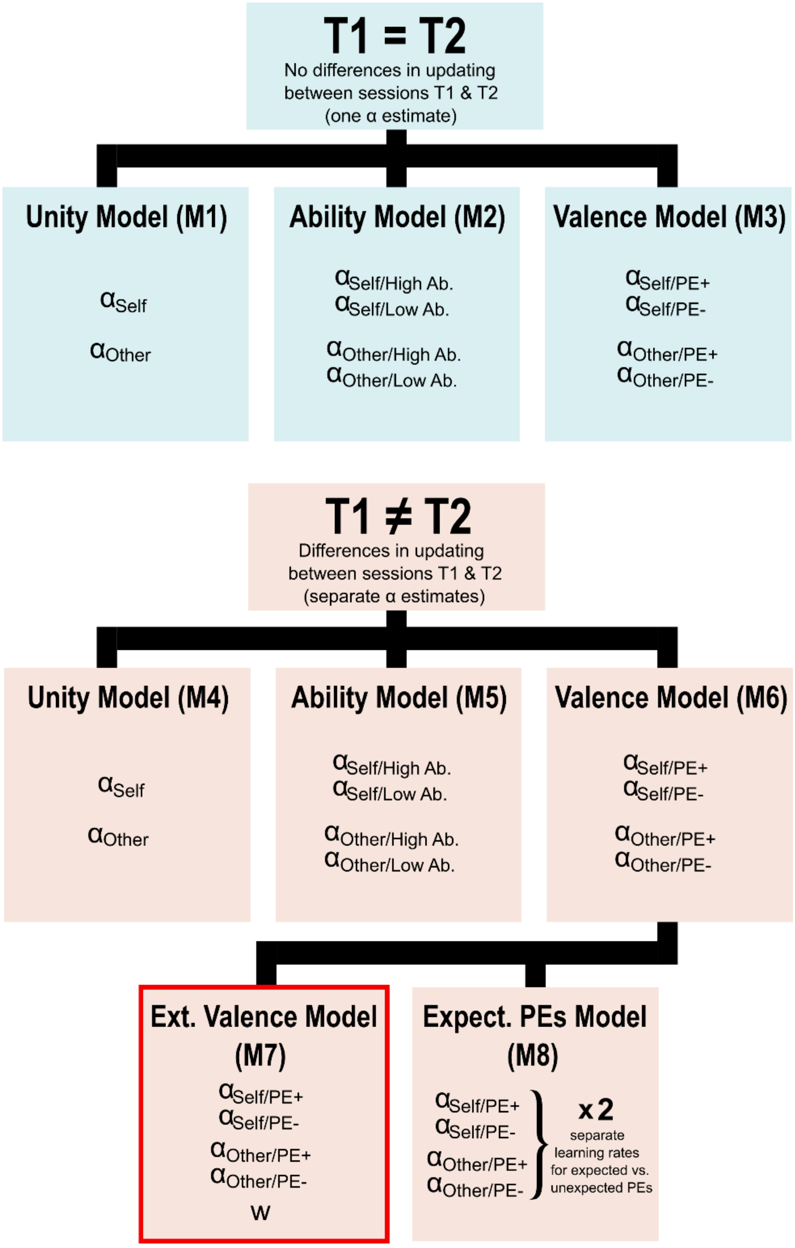


**All Models: starting values (initial expectation ratings) per category and session**

**(4 categories x 2 sessions)**

*Supplementary Figure 1*. Depiction of the model space. The first set of models (M1 to M3) assumed that the same learning rates were sufficient to accurately describe learning at both sessions (*T1=T2*). The second set (M4 to M8) assumed different learning rates for the two sessions (*T1≠T2*). Learning rates were segregated between the factors *Agent (Self* vs. *Other)* and the impact of prediction error valence (Valence Model/ Extended Valence Model; no impact: Unity Model) or *Ability* (Ability Model). A simple Mean Model (M9, not shown in the figure) with means for each condition was added to test whether stable assumptions in each *Ability* condition could explain expectation ratings better than PE learning. The Extended Valence Model (M7), which included a decay factor *w*, was the winning model for both sessions and also assumed different learning rates for *T1* and *T2*. Note that all models included estimates for initial expectations ratings for each category and each session (4 categories x 2 sessions) in addition to the depicted learning rates. See the Methods section for more information. $\alpha$ = learning rate, $w$ = weighting factor.

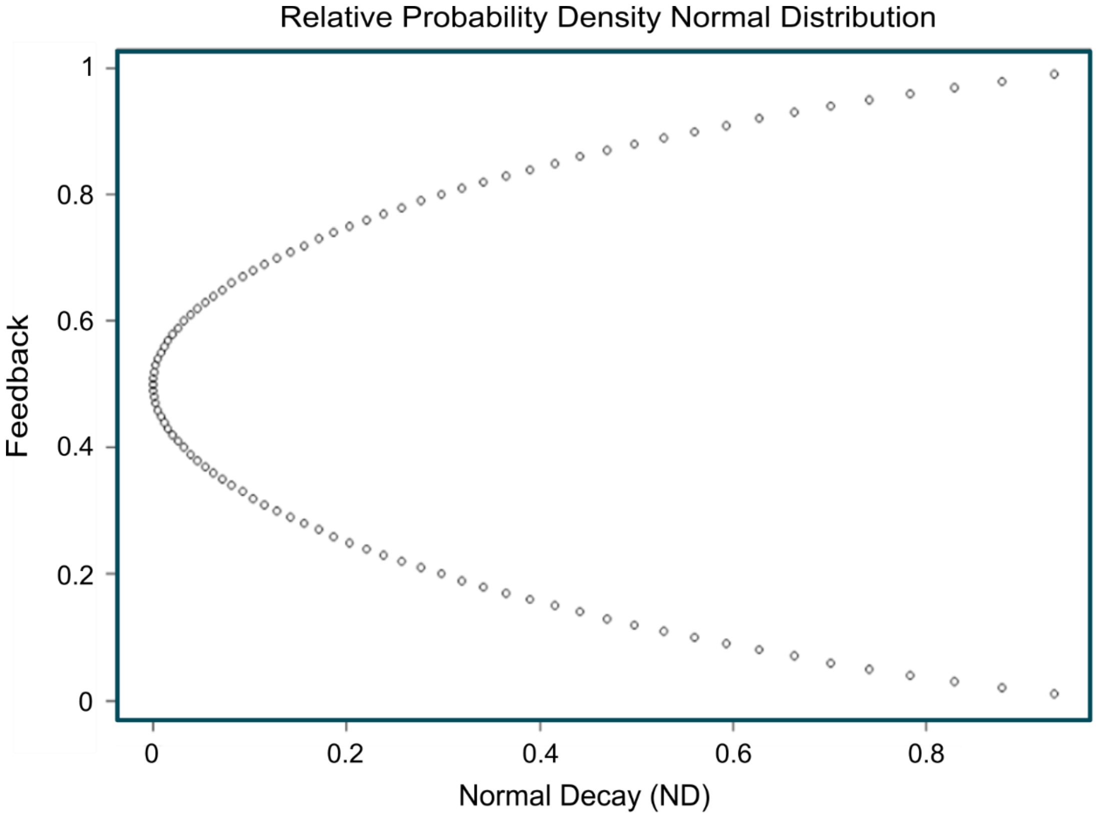


*Supplementary Figure 2.* Depiction of decay that follows the relative probability density of the normal distribution. Depicted values were introduced to the Extended Valence Model (M7) and were weighted by a weighting factor *w* (Müller-Pinzler et al., 2022). See the Methods section for more details.

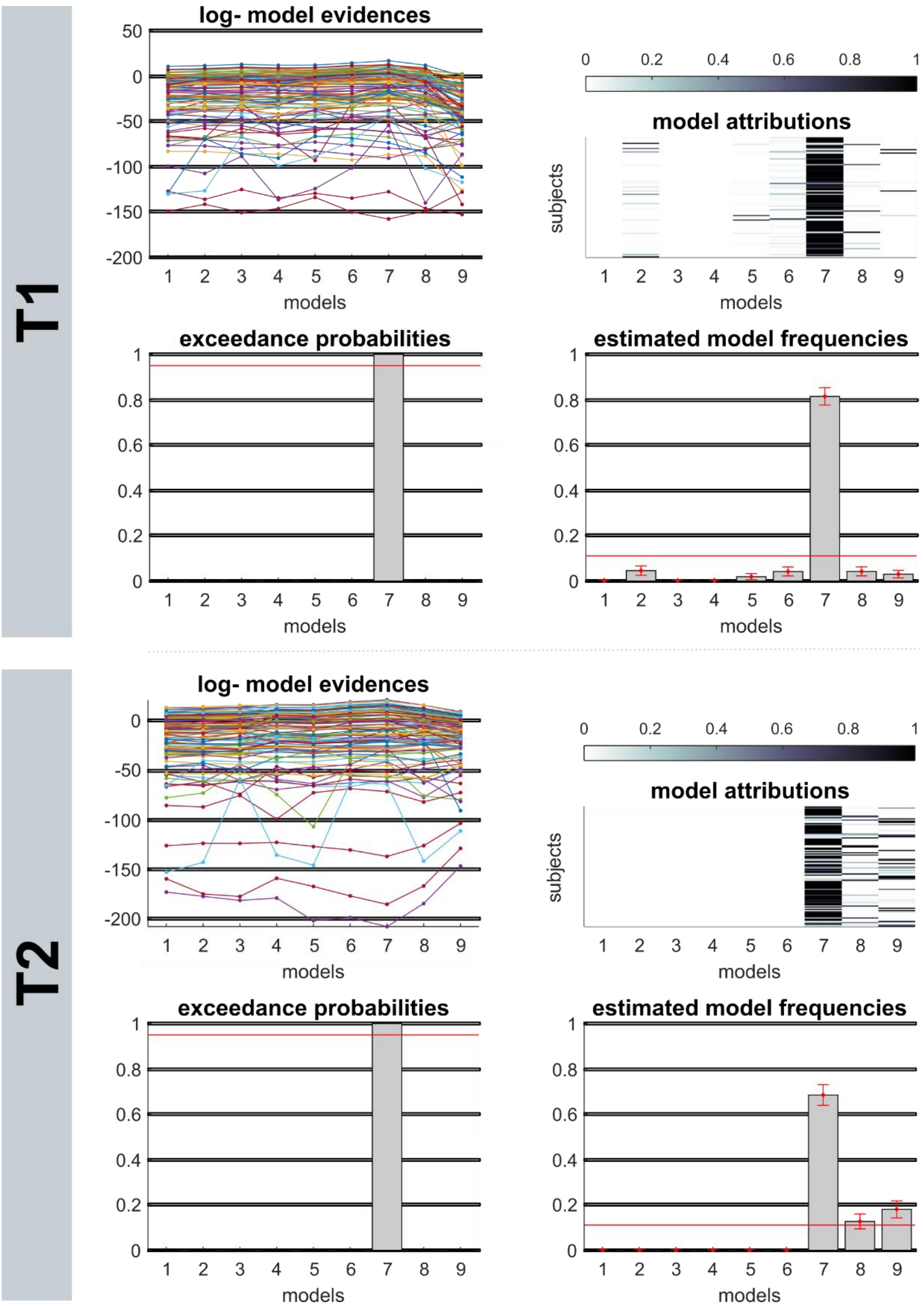


*Supplementary Figure 3*. Bayesian Model Selection (BMS). Results of BMS-process for *T1* and *T2*. Model 7 (Extended Valence Model) emerged as the winning model of both sessions with protected exceedance probabilities *pxp* > .999 (*T1* and *T2*) and an estimated model frequency of 81.98 for *T1* and 68.86 for *T2*.

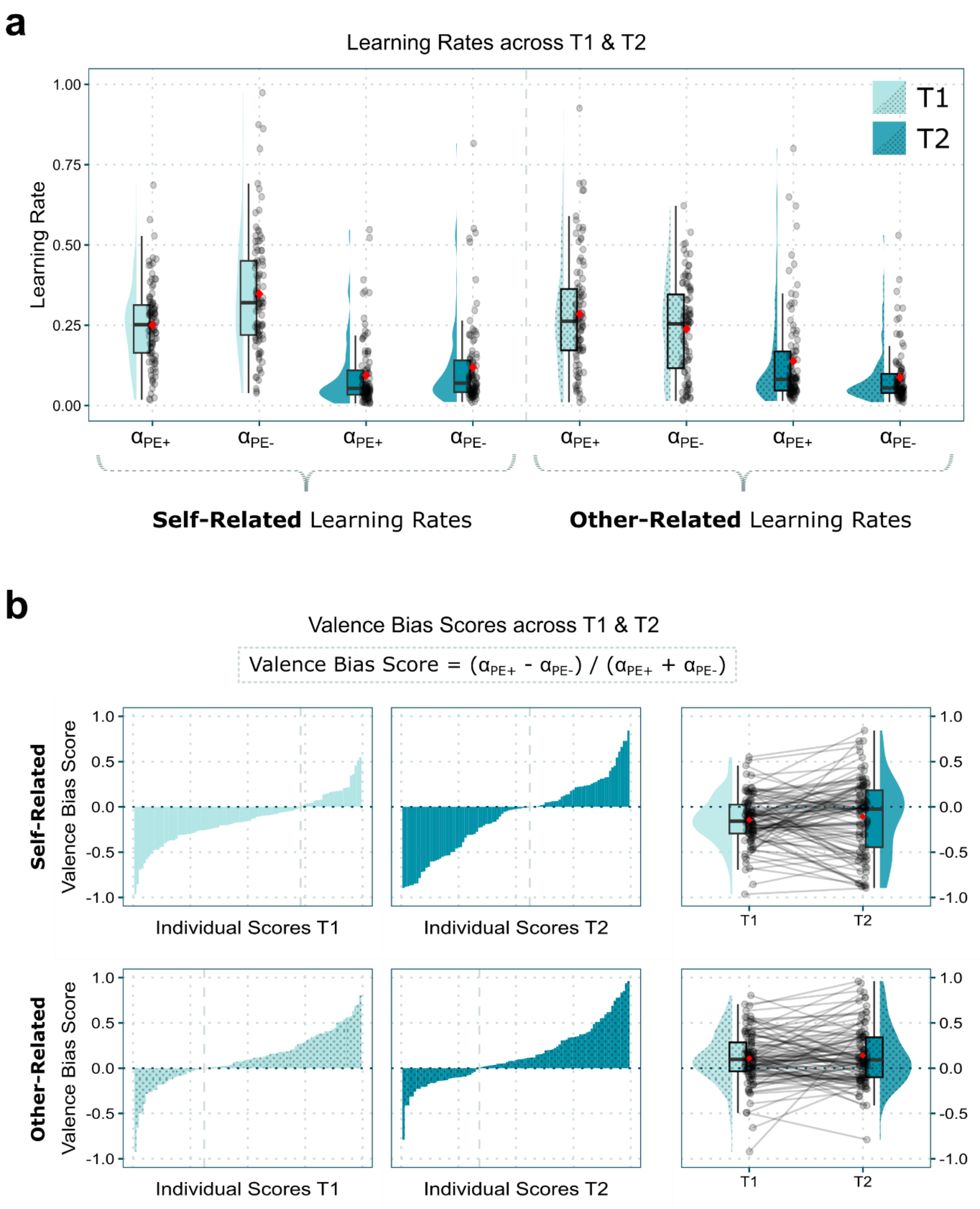


*Supplementary Figure 4*. Learning parameters for updating self- and other-related beliefs across the two sessions. **a.** Learning rates derived from the winning model for sessions *T1* and *T2*. **b.** Valence bias scores across the two sessions.

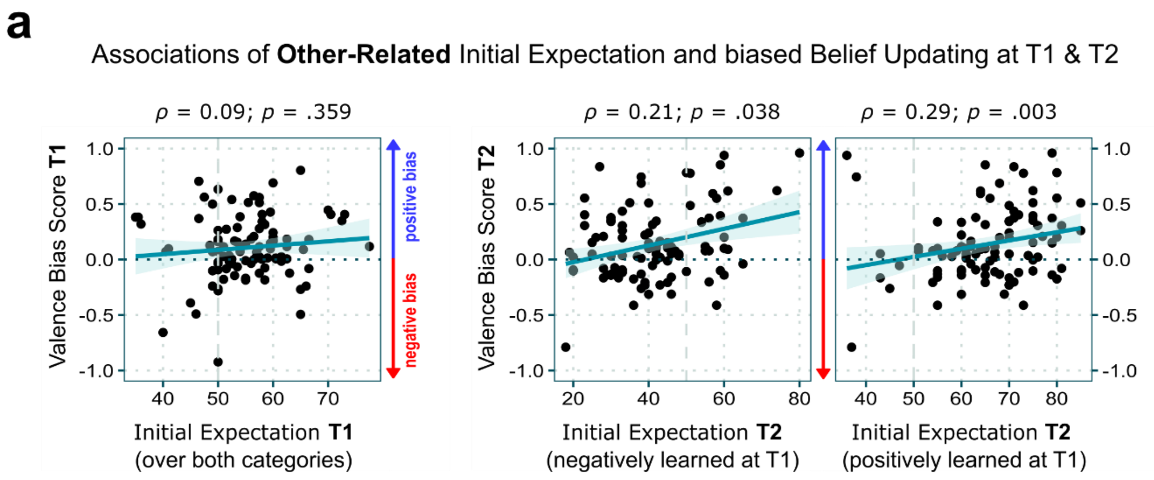


*Supplementary Figure 5.* Associations of other-related initial expectations and valence bias scores across the two sessions. $\rho$ = Spearman's rank correlation coefficient.

### Supplementary Notes

***Supplementary Note 1:*** Bayesian Model Selection.

Trial-by-trial changes of expectations within each condition (combinations of levels of the factors *Agent* and *Ability*) were modeled through updates from prediction errors (PEs). Learning rates were estimated in a way, that allowed for tests of PE-valence-related (positive vs. negative) and *Self* vs *Other*-related updating. In addition, models were considered that assumed equal vs. different learning rates for the two sessions *T1* vs. *T2*. Just like in our previous studies (Czekalla et al., 2021; Müller-Pinzler et al., 2022, 2019), the winning model was a valence-based model that included separate learning rates for positive vs. negative PEs and for *Self* vs. *Other* (see "Method" section for a more detailed explanation of the modeling process and the other models tested). For this study, this model additionally assumed different learning rates for the two sessions *T1* and *T2*. This Extended Valence Model, which also included a parameter *w* that modulates learning from more extreme feedback, received the highest sum PSIS-LOO scores out of all tested models (approximate leave-one-out cross-validation (LOO) using Pareto smoothed importance sampling (PSIS) (Vehtari et al., 2016); see *Supplementary Table 1* for all PSIS-LOO scores). For *T1*, Bayesian Model Selection (BMS) showed a protected exceedance probability of $pxp > .999$ for the extended Valence Model and a Bayesian Omnibus Risk of $BOR < .001$ with an expected model frequency of 81.89. For *T2*, the exceedance probability was $pxp > .999$ and the Bayesian Omnibus Risk was $BOR < .001$. Here, the expected model frequency was 68.86 (also see *Supplementary Figure 3* for plotted results of BMS). As a result of this selection process, all further analysis of learning parameters was based on the Extended Valence Model that allows for comparisons of PE-valence-based updating processes. For both sessions (*T1* and *T2*), the performance expectation ratings, predicted by the winning model, successfully captured the effects of the model-free analysis of the actual data (see *Supplementary Note 3*).

***Supplementary Note 2:*** Analysis of performance expectation ratings – Subjects adjust their expectations according to the provided feedback during belief formation at *T1* and show an impaired belief updating at *T2*.

*Model-free analysis T1.* An iterative modelbuilding approach (see "Methods" for details) resulted in a linear mixed-effects model that included Intercept, Agent, Ability, and Trial as both random and fixed effects. In addition, the two-way and one three-way interactions were included as fixed effects. The results are summarized in *Supplementary Table 4*. Most noteworthy, a significant main effect of *Ability* ($\beta = -7.81$, 95% *CI* [-10.19; -5.42], $t_{(135.4)} = -6.42$, $p < .001$) and interaction of *Ability* and *Trial* ($\beta = -1.39$, 95% *CI* [-1.47; -1.31], $t_{(7520)} = -33.32$, $p < .001$) showed that participants updated their performance expectations according to the provided feedback throughout the task. A significant main effect of *Agent* ($\beta = 6.01$, 95% *CI* [3.72; 8.29], $t_{(139.74)} = 5.15$, $p < .001$) showed a more positive evaluation of the other person's performance than of one's performance. Further, a significant interaction of *Agent* × *Ability* × *Trial* ($\beta = 0.31$, 95% *CI* [0.2; 0.43], $t_{(7520)} = 5.3$, $p < .001$) indicated differential learning patterns between the *Ability* contexts of *Self* and *Others* throughout the initial learning at *T1*.

To control for potential confounds resulting from the fact, that one group of participants completed the two sessions of the task on two consecutive days (therefore slept in between the sessions) while a second group completed the task on the same day (therefore stayed awake) for an additional test, the factor *Group (wake* vs. *sleep)* together with its two-way interactions was added to the final model. As it was expected for *T1* (where all measurements took place before potential sleep-related consolidation), no significant results for the factor *Group* could be found (main effect *Group*: $\beta = 2.92$, 95% *CI* [-1.99; 7.83], $t_{(97)} = 1.17$, $p = .247$; *Agent* × *Group*: $\beta = -3.84$, 95% *CI* [-8.39; 0.72], $t_{(97)} = -1.65$, $p = .102$; *Ability* ×*Group*: $\beta = -2.08$, 95% *CI* [-6.92; 2.76], $t_{(97)} = -0.84$, $p = .401$; *Trial* ×*Group*: $\beta = 0.03$, 95% *CI* [-0.11; 0.17], $t_{(97)} = 0.42$, $p = .676$).

*Model-free analysis T2.* For *T2* the above-mentioned modelbuilding approach was repeated and resulted in a linear mixed-effects model of identical structure. The summarized results are shown in *Supplementary Table 5.* Here, most importantly, a significant main effect of *Ability* ($\beta = 22.6$, 95% *CI* [19.45; 25.76], $t_{(115.6)} = 14.04$, $p < .001$) and interaction of *Ability* and *Trial* ($\beta = -0.25$, 95% *CI* [-0.33; -0.17], $t_{(7520)} = -6.23$, $p < .001$) showed tendencies of belief maintenances since performance expectation ratings tended to be higher in the estimation category of the *Low Ability* condition (the category that was positively learned at *T1*). Once again, as found at *T1*, subjects evaluated themselves more negatively than others (main effect *Agent*: $\beta = 8.79$, 95% *CI* [5.55; 12.04], $t_{(114.5)} = 5.31$, $p < .001$) and a significant interaction of *Agent* × *Ability* × *Trial* ($\beta = -0.12$, 95% *CI* [-0.23; 0.001], $t_{(7520)} = -2.03$, $p = .042$) showed differential learning patterns for *Self* and *Other* for the levels of *Ability* over the trials of *T2.*

Since potential effects of sleep-related consolidation could be assumed for *T2*, additional analysis with the added factor *Group* and its two-way interaction was conducted as it was done for session *T1*. Again, no significant effects were observed (main effect *Group*: $\beta = 2.34$, 95% *CI* [-2.98; 7.66], $t_{(97)} = 0.86$, $p = .391$; *Agent* × *Group*: $\beta = -5.37$, 95% *CI* [-12.18; 1.44], $t_{(97)} = -1.55$, $p = .125$; *Ability* ×*Group*: $\beta = 1.69$, 95% *CI* [-4.99; 8.38], $t_{(97)} = 0.50$, $p = .62$; *Trial* ×*Group*: $\beta = -0.01$, 95% *CI* [-0.12; 0.10], $t_{(97)} = -0.21$, $p = .835$).

***Supplementary Note 3:*** Posterior predictive checks: analyses of performance expectation ratings simulated by the winning model.

To assess whether the winning learning model could capture the effects of our model-free analyses (*Supplementary Note 1*), we let the winning model predict the time course of performance expectation ratings for each subject and repeated the analyses of the observed behavioral data on the simulated data. These posterior predictive checks followed the same procedure described in the methods section and *Supplementary Note 1.* The core effects of our observed data could successfully be captured. The summarized results are shown in *Supplementary Tables 6* and *7*. See *Figure 2* for a visual confirmation. Because the final models of the observed data included all main effects and interactions as fixed effects, the three-way interaction was still included for the posterior predictive check model of *T2*, although it did not add any further value to the final model ($X^2(1) = 1.32$, $p = .25$).

***Supplementary Note 4:*** Analysis of learning rates. Subjects show higher learning rates during the initial learning phase of *T1* compared to the revision phase of *T2*.

The iterative modelbuilding approach (see "Methods" for details) resulted in a linear mixed-effects model that included *Intercept, Agent, PE-Valence,* and *Session* as fixed effects and *Intercept, Agent,* and *Session* as random effects (inclusion of *PE-Valence* as a random effect caused issues with model convergence). In addition, the two-way-interactions and the one three-way-interaction were included as fixed effects in the final model. All results are summarized in *Supplementary Table 8.* As the most important findings of this analyses, firstly, the self-related negativity bias at *T1* found in previous studies (Czekalla et al., 2021; Müller-Pinzler et al., 2022, 2019) could be replicated (interaction *Agent* × *PE-Valence* × *Session:* $\beta = 0.07$, 95% *CI* [0.01; 0.13], $t_{(490)} = 2.15$, $p = .032$; pairwise comparisons of estimated marginal means: $\alpha_{Self/PE+/T1} - \alpha_{Self/PE-/T1}$: $t_{(490)} = -6.08$, $p_{Tukey} < .001$, $M_{Diff} = -0.1$, $SE = .02$; $\alpha_{Self/PE+/T2} - \alpha_{Self/PE-/T2}$: $t_{(490)} = -1.56$, $p_{Tukey} = .405$, $M_{Diff} = -0.02$, $SE = 0.02$). Secondly, overall learning rates at *T2* were significantly lower than at *T1* (main effect of *Session:* $\beta = -0.16$, 95% *CI* [-0.19; -0.12], $t_{(358.23)} = -8.23$, $p < .001$). That shows that subjects maintained their previously established beliefs rather than adjusted to the new feedback.

To once again rule out the potential effects of sleep-related processes of consolidation, the factor *Group (wake* vs. *sleep)* and its two-way interactions were added to the final model for additional analysis. No significant effects were observed (main effect *Group*: $\beta = 0.01$, 95% *CI* [-0.04; 0.06], $t_{(125.67)} = 0.34$, $p = .734$; *Agent* × *Group*: $\beta = 0.005$, 95% *CI* [-0.03; 0.04], $t_{(97)} = 0.25$, $p = .806$; *PE-Valence* × *Group*: $\beta = 0.003$, 95% *CI* [-0.03; 0.04], $t_{(489)} = 0.17$, $p = .868$; *Session* × *Group*: $\beta = -0.004$, 95% *CI* [-0.06; 0.05], $t_{(97)} = -0.14$, $p = .887$).

***Supplementary Note 5:*** Analysis of valence bias scores. Subjects show a negative bias in updating their self-beliefs and a positive bias when updating about the other person regardless of the session.

Modelbuilding (see “Methods” for details) resulted in a linear mixed-effects model that included *Intercept, Agent, Session* and the interaction of *Agent* and *Session* as fixed effects and the *Intercept* and *Agent* as random effects (inclusion of *Session* as random effect caused issues with convergence). The results are summarized in *Supplementary Table 9.* A significant main effect of *Agent* could be observed that showed a more positive updating regarding other-related beliefs (β = 0.25, 95% *CI* [0.17; 0.33], $t_{(98)} = 5.96$, $p < .001$). This was consistent across the two sessions (*T1* vs. *T2)*: No main effect for *Session* could be found (β = 0.03, 95% *CI* [-0.02; 0.08], $t_{(197)} = 1.35$, $p = .178$) and the interaction *Agent* × *Session* added no further value to the model ($X^2(1) < 0.01$, $p = .950$).

***Supplementary Note 6:*** Analysis of confidence ratings. Confidence grows during initial learning, regardless of one's ability. Conflicting feedback during the revision phase only results in a slight decrease.

An iterative modelbuilding approach (see "Methods" for details) resulted in a linear mixed-effects model that included *Intercept, Ability,* and *Timepoint* as fixed effects and *Intercept* as a random effect (addition of further random effects caused the model to not converge). The model did not benefit from the inclusion of the interaction of *Ability* and *Timepoint* ($X^2(3) = 1.56$, $p = .668$) as fixed effects, so the final model only considered the main effects as fixed effects. The final model showed no significant main effect of *Ability* (β = -1.03, 95% *CI* [-3.20; 1.14], $t_{(654)} = -0.93$, $p = .352$), suggesting that change of confidence did not depend on whether one expected to be good or less good in the respective domain. Most importantly, contrasts for the factor *Timepoint* showed, that confidence increased over the initial learning phase (*post T1* vs. *pre T1*: β = 11.15, 95% *CI* [8.09; 14.22], $t_{(654)} = 7.13$, $p < .001$) and, despite a small decrease, remained to be significantly higher than before the initial learning (*pre T2* vs. *pre T1*: β = 8.15, 95% *CI* [5.09; 11.22], $t_{(654)} = 5.21$, $p < .001$; *post T2* vs. *pre T1*: β = 6.64, 95% *CI* [3.57; 9.70], $t_{(654)} = 4.24$, $p < .001$). All results are also summarized in *Supplementary Table 10.*

As done for the analyses of expectation ratings and learning rates, an additional analysis was conducted by adding the factor *Group* and its two-way interactions to the final linear mixed-effects model for belief confidence. No significant main effect for *Group* was found (β = 4.2, 95% *CI* [-3.04; 11.43], $t_{(239.35)} = 1.14$, $p = .256$). While two *Timepoint* × *Group* contrasts slightly missed significance (*Timepoint pre T2* vs. *pre T1* × *Group*: β = 6.12, 95% *CI* [-0.49; 12.74], $t_{(650)} = 1.81$, $p = .07$; *Timepoint post T2* vs. *pre T1* × *Group*: β = 5.55, 95% *CI* [-1.06; 12.17], $t_{(650)} = 1.65$, $p = .1$) the contrast for ratings done before and after session *T1* reached significance (*Timepoint post T1* vs. *pre T1* × *Group*: β = 7.78, 95% *CI* [1.17; 14.40], $t_{(650)} = 2.31$, $p = .021$). However, since the confidence ratings of those time points were assessed before the actual sleep or wake phases, major confounds appear unlikely.

### Supplementary Tables

*Supplementary Table 1.* PSIS-LOO scores.

| Model | PSIS-LOO | LOO-SE | LOO-Diff | LOO-SE-Diff | % of $\hat{k}$ > 0.7 | No. Est. Parameters |
|---|---|---|---|---|---|---|
| **Session T1** | | | | | | |
| Mean Model (M9) | -3833.44 | 316.32 | -2035.37 | 175.09 | 0.01 | 8 |
| **T1 = T2** | | | | | | |
| Unity Model (M1) | -2570.13 | 322.26 | -772.06 | 125.56 | 0.40 | 10 |
| Ability Model (M2) | -2371.03 | 310.53 | -572.96 | 117.82 | 0.88 | 12 |
| Valence Model (M3) | -2191.13 | 294.13 | -393.07 | 62.78 | 0.38 | 12 |
| **T1 ≠ T2** | | | | | | |
| Unity Model (M4) | -2492.85 | 319.86 | -694.78 | 114.31 | 0.99 | 12 |
| Ability Model (M5) | -2305.30 | 305.72 | -507.23 | 112.56 | 1.58 | 16 |
| Valence Model (M6) | -2092.53 | 304.26 | -294.46 | 66.98 | 0.96 | 16 |
| Expected PEs Model (M8) | -2497.77 | 330.23 | -699.70 | 117.71 | 0.85 | 24 |
| Ext. Valence Model (M7) | -1798.07 | 298.87 | - | - | 1.54 | 18 |
| **Session T2** | | | | | | |
| Mean Model (M9) | -2146.34 | 296.19 | -548.28 | 159.99 | 0.01 | 8 |
| **T1 = T2** | | | | | | |
| Unity Model (M1) | -2255.90 | 347.91 | -657.84 | 120.19 | 0.21 | 10 |
| Ability Model (M2) | -2235.37 | 350.72 | -637.31 | 104.51 | 0.63 | 12 |
| Valence Model (M3) | -2083.07 | 333.26 | -485.01 | 66.01 | 0.28 | 12 |
| **T1 ≠ T2** | | | | | | |
| Unity Model (M4) | -1927.49 | 353.20 | -329.44 | 103.26 | 0.37 | 12 |
| Ability Model (M5) | -2038.22 | 374.42 | -440.16 | 111.10 | 1.09 | 16 |
| Valence Model (M6) | -1694.39 | 352.11 | -96.33 | 29.86 | 0.38 | 16 |
| Expected PEs Model (M8) | -1916.61 | 361.57 | -318.55 | 105.97 | 0.50 | 24 |
| Ext. Valence Model (M7) | -1598.06 | 366.95 | - | - | 0.39 | 18 |

*Note.* PSIS-LOO = sum score of approximate leave-one-out cross-validation (LOO) using pareto-smoothed importance sampling (PSIS); LOO-SE = standard error of PSIS-LOO; LOO-Diff = difference in expected predictive accuracy for all models in reference to the winning model with the highest PSIS-LOO (M7, Extended Valence Model) and standard error of these differences; percentage of $\hat{k}$ (estimated shape parameters of the generalized Pareto distribution) that exceed 0.7 (as suggested by Vehtari et al., 2016); No. Est. Parameters = number of parameters estimated by the model (learning rates and initial expectation ratings for each condition and session and weighting factor *w* in case of M7 (compare *Supplementary Figure 1*)). Note that the models were estimated for *T1* and *T2* together and, accordingly, the number of parameters for the estimation over both sessions is given in this column. Shaded rows mark the winning model per session.

*Supplementary Table 2*. Pre and post expectation ratings.

| Ability context (as assigned at T1) | Session T1 | | | | Session T2 | | | |
|---|---|---|---|---|---|---|---|---|
| | Pre T1 expectation | | Post T1 expectation | | Pre T2 expectation | | Post T2 expectation | |
| | *M* | *SD* | *M* | *SD* | *M* | *SD* | *M* | *SD* |
| **Self** | | | | | | | | |
| High Ability | 48.03 | 11.23 | 63.67 | 14.74 | 59.81 | 13.56 | 55.43 | 17.45 |
| Low Ability | 49.27 | 10.76 | 28.99 | 11.71 | 33.49 | 12.73 | 38.63 | 17.03 |
| **Other** | | | | | | | | |
| High Ability | 54.20 | 9.62 | 69.38 | 10.80 | 65.61 | 10.48 | 62.08 | 13.69 |
| Low Ability | 56.46 | 8.17 | 39.68 | 14.98 | 41.69 | 12.96 | 50.88 | 15.60 |

Note. *M* = mean; *SD* = standard deviation. Pre (first trial expectation rating) and post (last trial expectation rating) expectation ratings for each of the four *Ability* conditions per session (*T1* and *T2*).

*Supplementary Table 3*. Pre and post confidence ratings.

| Ability context (as assigned at T1) | Session T1 | | | | Session T2 | | | |
|---|---|---|---|---|---|---|---|---|
| | Pre T1 confidence | | Post T1 confidence | | Pre T2 confidence | | Post T2 confidence | |
| | *M* | *SD* | *M* | *SD* | *M* | *SD* | *M* | *SD* |
| **Self** | | | | | | | | |
| High Ability | 52.26 | 27.89 | 65.44 | 15.37 | 62.02 | 14.26 | 59.39 | 16.70 |
| Low Ability | 53.21 | 26.67 | 62.90 | 16.23 | 60.26 | 14.40 | 59.06 | 16.06 |
| **Other** | | | | | | | | |
| High Ability | 43.04 | 28.44 | 67.76 | 14.16 | 63.72 | 15.43 | 61.84 | 17.33 |
| Low Ability | 40.96 | 27.49 | 60.63 | 14.27 | 58.72 | 15.82 | 59.28 | 16.52 |

Note. *M* = mean; *SD* = standard deviation. Pre and post confidence ratings for each of the four *Ability* conditions per session (*T1* and *T2*).

*Supplementary Table 4*. Model comparison of linear mixed-effects models on performance expectation ratings at *T1* and results of the final model.

| | Model fit | | | Chi-square test against nested | | |
|---|---|---|---|---|---|---|
| **Added fixed effects** | AIC | BIC | LL | df | $X^2$ | *p* |
| Intercept | 57593.68 | 57677.41 | -28784.84 | | | |
| Agent | 57574.32 | 57665.02 | -28774.16 | 1 | 21.37 | <.001*** |
| Ability | 57423.62 | 57521.30 | -28697.81 | 1 | 152.70 | <.001*** |
| Trial | 57421.77 | 57526.43 | -28695.88 | 1 | 3.85 | .05* |
| Agent × Ability | 57404.67 | 57516.31 | -28686.34 | 1 | 19.09 | <.001*** |
| Ability × Trial | 55846.66 | 55965.27 | -27906.33 | 1 | 1560.02 | <.001*** |
| Agent × Trial | 55802.85 | 55928.44 | -27883.43 | 1 | 45.80 | <.001*** |
| Agent × Ability × Trial | 55776.83 | 55909.39 | -27869.41 | 1 | 28.03 | <.001*** |

**Estimates of the final model**

| | | | 95% CI | | | |
|---|---|---|---|---|---|---|
| **Fixed effects** | *β* | *SE* | lower | upper | *t* | *p* |
| Intercept | 52.23 | 1.18 | 49.93 | 54.54 | 44.37 | <.001*** |
| Agent | 6.01 | 1.17 | 3.72 | 8.29 | 5.15 | <.001*** |
| Ability | -7.81 | 1.22 | -10.19 | -5.42 | -6.42 | <.001*** |
| Trial | 0.53 | 0.04 | 0.45 | 0.61 | 12.97 | <.001*** |
| Agent × Ability | -1.63 | 0.71 | -3.01 | -0.24 | -2.30 | .022* |
| Ability × Trial | -1.39 | 0.04 | -1.47 | -1.31 | -33.32 | <.001*** |
| Agent × Trial | 0.04 | 0.04 | -0.04 | 0.13 | 1.05 | .292 |
| Agent × Ability × Trial | 0.31 | 0.06 | 0.20 | 0.43 | 5.30 | <.001*** |

| **Random effects** | *Variance* | *SD* |
|---|---|---|
| Participant (Intercept) | 124.82 | 11.17 |
| Agent | 109.74 | 10.48 |
| Ability | 121.78 | 11.04 |
| Trial | 0.08 | 0.28 |
| **Model fit** | Marginal $R^2$ | Conditional $R^2$ |
| | 0.51 | 0.8 |

*Note*. *AIC* = Akaike information criterion; *BIC* = Bayesian information criterion; *LL* = log-likelihood; *df* = degrees of freedom; *β* = regression coefficient; *SE* = standard error of the mean; 95% CI = 95% confidence intervals of regression coefficient estimates; *SD* = standard deviation. The shaded row marks the final model chosen for the analysis. Equation of final model: expectation rating *T1* ~ (agent + ability + trial + 1|subject) + agent + ability + trial + agent*ability + ability*trial + agent*trial + agent*ability*trial. *p*-values for fixed effects were calculated using Satterthwaite's approach. * $p < .05$, ** $p < .01$, *** $p < .001$.

*Supplementary Table 5*. Model comparison of linear mixed-effects models on performance expectation ratings at *T2* and results of the final model.

| | Model fit | | | Chi-square test against nested | | |
|---|---|---|---|---|---|---|
| **Added fixed effects** | AIC | BIC | LL | df | $X^2$ | *p* |
| Intercept | 55736.11 | 55819.84 | -27856.06 | | | |
| Agent | 55712.73 | 55803.43 | -27843.36 | 1 | 25.38 | <.001*** |
| Ability | 55629.25 | 55726.93 | -27800.63 | 1 | 85.48 | <.001*** |
| Trial | 55629.50 | 55734.16 | -27799.75 | 1 | 1.75 | .186 |
| Agent × Ability | 55492.10 | 55603.73 | -27730.05 | 1 | 139.40 | <.001*** |
| Ability × Trial | 55377.62 | 55496.23 | -27671.81 | 1 | 116.48 | <.001*** |
| Agent × Trial | 55357.96 | 55483.55 | -27660.98 | 1 | 21.66 | <.001*** |
| Agent × Ability × Trial | 55355.82 | 55488.39 | -27658.91 | 1 | 4.13 | .042* |

**Estimates of the final model**

| | | | 95% CI | | | |
|---|---|---|---|---|---|---|
| **Fixed effects** | *β* | *SE* | lower | upper | *t* | *p* |
| Intercept | 35.51 | 1.26 | 33.03 | 37.99 | 28.10 | <.001*** |
| Agent | 8.79 | 1.66 | 5.55 | 12.04 | 5.31 | <.001*** |
| Ability | 22.60 | 1.61 | 19.45 | 25.76 | 14.04 | <.001*** |
| Trial | 0.09 | 0.03 | 0.02 | 0.16 | 2.63 | .009** |
| Agent × Ability | -2.71 | 0.68 | -4.05 | -1.38 | -3.98 | <.001*** |
| Ability × Trial | -0.25 | 0.04 | -0.33 | -0.17 | -6.23 | <.001*** |
| Agent × Trial | 0.19 | 0.04 | 0.11 | 0.27 | 4.73 | <.001*** |
| Agent × Ability × Trial | -0.12 | 0.06 | -0.23 | 0.00 | -2.03 | .042* |

| **Random effects** | *Variance* | *SD* |
|---|---|---|
| Participant (Intercept) | 146.57 | 12.11 |
| Agent | 248.29 | 15.76 |
| Ability | 233.71 | 15.29 |
| Trial | 0.04 | 0.20 |

| **Model fit** | Marginal $R^2$ | Conditional $R^2$ |
|---|---|---|
| | 0.33 | 0.83 |

*Note. AIC* = Akaike information criterion; *BIC* = Bayesian information criterion; *LL* = log-likelihood; *df* = degrees of freedom; *β* = regression coefficient; *SE* = standard error of the mean; 95% CI = 95% confidence intervals of regression coefficient estimates; *SD* = standard deviation. The shaded row marks the final model chosen for the analysis. Equation of final model: expectation rating *T2* ~ (agent + ability + trial + 1|subject) + agent + ability + trial + agent*ability + ability*trial + agent*trial + agent*ability*trial. *p*-values for fixed effects were calculated using Satterthwaite's approach. * $p < .05$, ** $p < .01$, *** $p < .001$.

*Supplementary Table 6.* Posterior predictive check of simulated data: Model comparison of linear mixed-effects models on simulated performance expectation ratings at *T1* and final model results.

| | Model fit | | | Chi-square test against nested | | |
|---|---|---|---|---|---|---|
| **Added fixed effects** | AIC | BIC | LL | df | $X^2$ | *p* |
| Intercept | 53905.01 | 53988.74 | -26940.50 | | | |
| Agent | 53887.75 | 53978.45 | -26930.87 | 1 | 19.26 | <.001*** |
| Ability | 53723.52 | 53821.20 | -26847.76 | 1 | 166.23 | <.001*** |
| Trial | 53719.95 | 53824.60 | -26844.97 | 1 | 5.57 | .018* |
| Agent × Ability | 53703.79 | 53815.43 | -26835.90 | 1 | 18.15 | <.001*** |
| Ability × Trial | 50384.22 | 50502.83 | -25175.11 | 1 | 3321.57 | <.001*** |
| Agent × Trial | 50294.88 | 50420.47 | -25129.44 | 1 | 91.35 | <.001*** |
| Agent × Ability × Trial | 50285.69 | 50418.25 | -25123.84 | 1 | 11.19 | <.001*** |

**Estimates of the final model**

| | | | 95% CI | | | |
|---|---|---|---|---|---|---|
| **Fixed effects** | $\beta$ | *SE* | lower | upper | *t* | *p* |
| Intercept | 51.91 | 1.12 | 49.73 | 54.10 | 46.48 | <.001*** |
| Agent | 5.41 | 1.11 | 3.25 | 7.58 | 4.90 | <.001*** |
| Ability | -7.06 | 1.05 | -9.12 | -4.99 | -6.70 | <.001*** |
| Trial | 0.53 | 0.04 | 0.46 | 0.60 | 14.41 | <.001*** |
| Agent × Ability | -0.18 | 0.49 | -1.14 | 0.79 | -0.36 | .722 |
| Ability × Trial | -1.40 | 0.03 | -1.46 | -1.35 | -48.36 | <.001*** |
| Agent × Trial | 0.13 | 0.03 | 0.07 | 0.19 | 4.42 | <.001*** |
| Agent × Ability × Trial | 0.14 | 0.04 | 0.06 | 0.22 | 3.35 | <.001*** |

| **Random effects** | *Variance* | *SD* |
|---|---|---|
| Participant (Intercept) | 117.54 | 10.84 |
| Agent | 108.91 | 10.44 |
| Ability | 97.66 | 9.88 |
| Trial | 0.09 | 0.30 |

| **Model fit** | Marginal $R^2$ | Conditional $R^2$ |
|---|---|---|
| | 0.58 | 0.89 |

*Note. AIC* = Akaike information criterion; *BIC* = Bayesian information criterion; *LL* = log-likelihood; *df* = degrees of freedom; $\beta$ = regression coefficient; *SE* = standard error of the mean; 95% CI = 95% confidence intervals of regression coefficient estimates; *SD* = standard deviation. The shaded row marks the final model chosen for the analysis. Equation of final model: predicted expectation rating *T1* ~ (agent + ability + trial + 1|subject) + agent + ability + trial + agent*ability + ability*trial + agent*trial + agent*ability*trial. *p*-values for fixed effects were calculated using Satterthwaite's approach. * $p < .05$, ** $p < .01$, *** $p < .001$.

*Supplementary Table 7.* Posterior predictive check of simulated data: Model comparison of linear mixed-effects models on simulated performance expectation ratings at *T2* and final model results.

| | Model fit | | | Chi-square test against nested | | |
|---|---|---|---|---|---|---|
| **Added fixed effects** | AIC | BIC | LL | df | $X^2$ | *p* |
| Intercept | 50486.48 | 50570.20 | -25231.24 | | | |
| Agent | 50462.40 | 50553.10 | -25218.20 | 1 | 26.08 | <.001*** |
| Ability | 50383.95 | 50481.63 | -25177.98 | 1 | 80.45 | <.001*** |
| Trial | 50383.32 | 50487.97 | -25176.66 | 1 | 2.63 | .105 |
| Agent × Ability | 50095.68 | 50207.31 | -25031.84 | 1 | 289.64 | <.001*** |
| Ability × Trial | 49055.50 | 49174.12 | -24510.75 | 1 | 1042.17 | <.001*** |
| Agent × Trial | 49001.63 | 49127.22 | -24482.82 | 1 | 55.87 | <.001*** |
| Agent × Ability × Trial | 49002.31 | 49134.88 | -24482.16 | 1 | 1.32 | .25 |

**Estimates of the final model**

| | | | 95% CI | | | |
|---|---|---|---|---|---|---|
| **Fixed effects** | *β* | *SE* | lower | upper | *t* | *p* |
| Intercept | 33.87 | 1.25 | 31.42 | 36.31 | 27.13 | <.001*** |
| Agent | 9.31 | 1.63 | 6.12 | 12.51 | 5.71 | <.001*** |
| Ability | 25.59 | 1.58 | 22.50 | 28.68 | 16.24 | <.001*** |
| Trial | 0.27 | 0.03 | 0.21 | 0.33 | 9.33 | <.001*** |
| Agent × Ability | -3.52 | 0.45 | -4.40 | -2.65 | -7.89 | <.001*** |
| Ability × Trial | -0.60 | 0.03 | -0.66 | -0.55 | -22.91 | <.001*** |
| Agent × Trial | 0.16 | 0.03 | 0.11 | 0.21 | 6.11 | <.001*** |
| Agent × Ability × Trial | -0.04 | 0.04 | -0.12 | 0.03 | -1.15 | .25 |

| **Random effects** | *Variance* | *SD* |
|---|---|---|
| Participant (Intercept) | 149.37 | 12.22 |
| Agent | 253.86 | 15.93 |
| Ability | 235.95 | 15.36 |
| Trial | 0.05 | 0.22 |
| **Model fit** | Marginal $R^2$ | Conditional $R^2$ |
| | 0.35 | 0.92 |

*Note. AIC* = Akaike information criterion; *BIC* = Bayesian information criterion; *LL* = log-likelihood; *df* = degrees of freedom; *β* = regression coefficient; *SE* = standard error of the mean; 95% CI = 95% confidence intervals of regression coefficient estimates; *SD* = standard deviation. The shaded row marks the final model chosen for the analysis. Note, that in this case the model including the three-way interaction was chosen although this model was not the clear winner. This was done to ensure a better comparison with the model used to analyze the observed data (see *Supplementary Table 5*). Equation of final model: predicted expectation rating *T2* ~ (agent + ability + trial + 1|subject) + agent + ability + trial + agent*ability + ability*trial + agent*trial + agent*ability*trial. *p*-values for fixed effects were calculated using Satterthwaite's approach. * $p < .05$, ** $p < .01$, *** $p < .001$.

*Supplementary Table 8.* Model comparison of linear mixed-effects models on learning rates over both sessions (*T1* and *T2*) and final model results.

| | Model fit | | | Chi-square test against nested | | |
|---|---|---|---|---|---|---|
| **Added fixed effects** | AIC | BIC | LL | df | $X^2$ | *p* |
| Intercept | -794.5440 | -757.1476 | 405.2720 | | | |
| Agent | -800.6782 | -758.6072 | 409.3391 | 1 | 8.13 | .004** |
| PE-Valence | -799.2674 | -752.5217 | 409.6337 | 1 | 0.59 | .443 |
| Session | -898.0992 | -846.6790 | 460.0496 | 1 | 100.83 | <.001*** |
| Agent × PE-Valence | -939.7252 | -883.6305 | 481.8626 | 1 | 43.63 | <.001*** |
| Agent × Session | -945.1060 | -884.3367 | 485.5530 | 1 | 7.38 | .007** |
| PE-Valence × Session | -948.7471 | -883.3032 | 488.3735 | 1 | 5.64 | .018* |
| Agent × PE-Valence × Session | -951.3790 | -881.2606 | 490.6895 | 1 | 4.63 | .031* |

**Estimates of the final model**

| | | | 95% CI | | | |
|---|---|---|---|---|---|---|
| **Fixed effects** | *β* | *SE* | lower | upper | *t* | *p* |
| Intercept | 0.25 | 0.01 | 0.22 | 0.28 | 17.97 | <.001*** |
| Agent | 0.03 | 0.02 | 0.00 | 0.07 | 2.09 | .037* |
| PE-Valence | 0.10 | 0.02 | 0.07 | 0.13 | 6.08 | <.001*** |
| Session | -0.16 | 0.02 | -0.19 | -0.12 | -8.23 | <.001*** |
| Agent × PE-Valence | -0.14 | 0.02 | -0.19 | -0.10 | -6.35 | <.001*** |
| Agent × Session | 0.01 | 0.02 | -0.03 | 0.05 | 0.42 | .674 |
| PE-Valence × Session | -0.07 | 0.02 | -0.12 | -0.03 | -3.20 | .001*** |
| Agent × PE-Valence × Session | 0.07 | 0.03 | 0.01 | 0.13 | 2.15 | .032* |

| **Random effects** | *Variance* | *SD* |
|---|---|---|
| Participant (Intercept) | 0.006 | 0.081 |
| Session | 0.010 | 0.101 |
| Agent | 0.001 | 0.034 |

| **Model fit** | Marginal $R^2$ | Conditional $R^2$ |
|---|---|---|
| | 0.29 | 0.55 |

*Note. AIC* = Akaike information criterion; *BIC* = Bayesian information criterion; *LL* = log-likelihood; *df* = degrees of freedom; *β* = regression coefficient; *SE* = standard error of the mean; 95% CI = 95% confidence intervals of regression coefficient estimates; *SD* = standard deviation. The shaded row marks the final model chosen for the analysis. Equation of final model: learning rates ~ (session + agent + 1|subject) + agent + PE-valence + session + agent*PE-valence + agent*session + PE-valence*session + agent*PE-Valence*session. *p*-values for fixed effects were calculated using Satterthwaite's approach. * $p < .05$, ** $p < .01$, *** $p < .001$.

*Supplementary Table 9.* Model comparison of linear mixed-effects models on valence bias scores over both sessions (*T1* and *T2*) and final model results.

| | Model fit | | | Chi-square test against nested | | |
|---|---|---|---|---|---|---|
| **Added fixed effects** | AIC | BIC | LL | df | $X^2$ | *p* |
| Intercept | 233.9907 | 253.7877 | -111.94033 | | | |
| Agent | 205.2738 | 229.1623 | -96.63692 | 1 | 30.61 | <.001*** |
| Session | 205.4488 | 233.3287 | -95.72440 | 1 | 1.83 | .177 |
| Agent × Session | 207.4449 | 239.2962 | -95.72244 | 1 | < 0.01 | .950 |

**Estimates of the final model**

| | | | 95% CI | | | |
|---|---|---|---|---|---|---|
| **Fixed effects** | *β* | *SE* | lower | upper | *t* | *p* |
| Intercept | -0.14 | 0.03 | -0.21 | -0.08 | -4.49 | <.001*** |
| Agent | 0.25 | 0.04 | -0.17 | 0.33 | 5.96 | <.001*** |
| Session | 0.03 | 0.02 | -0.02 | 0.08 | 1.25 | .178 |

| **Random effects** | *Variance* | *SD* |
|---|---|---|
| Participant (Intercept) | 0.05 | 0.23 |
| Agent | 0.11 | 0.34 |

| **Model fit** | Marginal $R^2$ | Conditional $R^2$ |
|---|---|---|
| | 0.13 | 0.5 |

*Note. AIC* = Akaike information criterion; *BIC* = Bayesian information criterion; *LL* = log-likelihood; *df* = degrees of freedom; *β* = regression coefficient; *SE* = standard error of the mean; 95% CI = 95% confidence intervals of regression coefficient estimates; *SD* = standard deviation. The shaded row marks the final model chosen for the analysis. Note, that in this case the model including the fixed effect for the session was chosen although this model was not the clear winner. This was done to explicitly emphasize that the biases between the two sessions are comparable. Equation of final model: valence bias score ~ (agent + 1|subject) + agent + session. *p*-values for fixed effects were calculated using Satterthwaite's approach. * $p < .05$, ** $p < .01$, *** $p < .001$.

*Supplementary Table 10*. Model comparison of linear mixed-effects models on self-related confidence ratings over both sessions (*T1* and *T2*) and final model results.

| **Added fixed effects** | **Model fit** AIC | BIC | LL | **Chi-square test against nested** df | $X^2$ | *p* |
|---|---|---|---|---|---|---|
| Intercept | 6444.271 | 6458.139 | -3219.135 | | | |
| Ability | 6445.464 | 6463.955 | -3218.732 | 1 | 0.81 | .369 |
| Timepoint | 6398.788 | 6431.147 | -3192.394 | 3 | 52.68 | <.001*** |
| Ability × Timepoint | 6403.226 | 6449.453 | -3191.613 | 3 | 1.56 | .668 |

**Estimates of the final model**

| **Fixed effects** | *β* | *SE* | 95% CI lower | 95% CI upper | *t* | *p* |
|---|---|---|---|---|---|---|
| Intercept | 53.25 | 1.74 | 49.84 | 56.66 | 30.63 | <.001*** |
| Ability | -1.03 | 1.11 | -3.20 | 1.14 | -0.93 | .352 |
| Timepoint | | | | | | |
| *post T1* vs. *pre T1* | 11.15 | 1.56 | 8.09 | 14.22 | 7.13 | <.001*** |
| *pre T2* vs. *pre T1* | 8.15 | 1.56 | 5.09 | 11.22 | 5.21 | <.001*** |
| *post T2* vs. *pre T1* | 6.64 | 1.56 | 3.57 | 9.70 | 4.24 | <.001*** |

| **Random effects** | *Variance* | *SD* |
|---|---|---|
| Participant (Intercept) | 140.41 | 11.85 |

| **Model fit** | Marginal $R^2$ | Conditional $R^2$ |
|---|---|---|
| | 0.04 | 0.41 |

*Note*. *AIC* = Akaike information criterion; *BIC* = Bayesian information criterion; *LL* = log-likelihood; *df* = degrees of freedom; *β* = regression coefficient; *SE* = standard error of the mean; 95% CI = 95% confidence intervals of regression coefficient estimates; *SD* = standard deviation. The shaded row marks the final model chosen for the analysis. Equation of final model: confidence rating ~ (1|subject) + ability + timepoint. *p*-values for fixed effects were calculated using Satterthwaite's approach. * $p < .05$, ** $p < .01$, *** $p < .001$.

*Supplementary Table 11*. Distribution of positive and negative prediction errors across conditions.

| Condition (Agent × PE-Valence) | | *M* | *SD* | *M* frequency |
|---|---|---|---|---|
| | | **Session T1** | | |
| **Self** | Positive PEs | 13.35 | 1.17 | 20.79 |
| | Negative PEs | -12.38 | 1.67 | 19.21 |
| **Other** | Positive PEs | 13.05 | 1.44 | 19.03 |
| | Negative PEs | -13.31 | 1.38 | 20.97 |
| | | **Session T2** | | |
| **Self** | Positive PEs | 13.57 | 1.38 | 19.90 |
| | Negative PEs | -12.74 | 1.55 | 20.10 |
| **Other** | Positive PEs | 13.29 | 1.22 | 19.43 |
| | Negative PEs | -13.15 | 1.03 | 20.57 |

*Note*. PE = prediction error; *M* = mean; *SD* = standard deviation. Depicted are mean values and standard deviations of averaged PEs per subject. *M* frequency shows frequencies of PEs across conditions.
